# *Pseudomonas aeruginosa* AlgF is a protein-protein interaction mediator required for acetylation of the alginate exopolysaccharide

**DOI:** 10.1101/2023.07.25.550544

**Authors:** Kristin E. Low, Andreea A. Gheorghita, Stephanie D. Tammam, Gregory B. Whitfield, Yancheng E. Li, Laura M. Riley, Joel T. Weadge, Shane J. Caldwell, P. Andrew Chong, Marthe T. C. Walvoort, Elena N. Kitova, John S. Klassen, Jeroen D. C. Codée, P. Lynne Howell

**Author notes:** To whom correspondence should be addressed: P. Lynne Howell. These authors contributed equally to this work (Kristin E. Low, Andreea A. Gheorghita, Stephanie D. Tammam, Gregory B. Whitfield). K.E.L: Lethbridge Research and Development Centre, Agriculture and Agri-Food Canada, Lethbridge, Alberta, Canada; A.A.G.: Research Institute of Molecular Pathology, Vienna Biocenter, 1030 Vienna, Austria; G.B.W.: Département de Microbiologie, Infectiologie, et Immunologie, Université de Montréal, Montréal, Québec, Canada; Y.E.L.: Division of Chemistry and Chemical Engineering, California Institute of Technology, Pasadena, California, USA; L.M.R.: Ontario Genomics, Toronto, Ontario, Canada; J.T.W.: Department of Biology, Wilfrid Laurier University, Waterloo, Ontario, Canada; S.J.C: Amgen British Columbia, Burnaby, British Columbia, Canada; M.T.C.W: Stratingh Institute for Chemistry, University of Groningen, Groningen, The Netherlands.

## Abstract

Enzymatic modifications of bacterial exopolysaccharides enhance immune evasion and persistence during infection. In the Gram-negative opportunistic pathogen *Pseudomonas aeruginosa*, acetylation of alginate reduces opsonic killing by phagocytes and improves reactive oxygen species scavenging. Although it is well-known that alginate acetylation in *P. aeruginosa* requires AlgI, AlgJ, AlgF, and AlgX, how these proteins coordinate polymer modification at a molecular level remains unclear. Here, we describe the structural characterization of AlgF and its protein interaction network. We characterize direct interactions between AlgF and both AlgJ and AlgX *in vitro*, and demonstrate an association between AlgF and AlgX, as well as AlgJ and AlgI, in *P. aeruginosa*. We determine that AlgF does not exhibit acetylesterase activity and is unable to bind to polymannuronate *in vitro.* Therefore, we propose that AlgF functions to mediate protein-protein interactions between alginate acetylation enzymes, forming the periplasmic AlgJFXK (AlgJ-AlgF-AlgX-AlgK) acetylation and export complex required for robust biofilm formation.

## INTRODUCTION

Biofilms are communities of bacterial cells surrounded and protected by a self-produced matrix containing lipids, exopolysaccharides, extracellular DNA, and proteins, and more complex structures such as membrane vesicles, bacteriophage, and amyloid fibres (1–3). The biofilm matrix promotes adhesion and cohesion of bacterial cells, and permits bacteria to adapt and thrive as a multicellular community despite environmental stresses (2, 4, 5). Bacterial biofilms can form in a variety of environments, including niches relevant to human health. They can grow on solid surfaces (e.g., medical devices), at an air-liquid interface (e.g., dental biofilms), or within semi-solid media (e.g., sputum in the lungs of individuals with cystic fibrosis (CF) during chronic infection with *Pseudomonas aeruginosa*) (2).

Modification of exopolysaccharides within the biofilm confers protection to pathogenic bacteria during infection (6). Deacetylation of poly-*N*-acetyl-glucosamine (PNAG) is required for biofilm formation in *Streptococcus epidermidis*, *Streptococcus aureus*, *Escherichia coli*, and *Yersinia pestis*, and provides resistance to neutrophil phagocytosis and enhances persistence in a mouse model of infection for *S. epidermidis* and *S. aureus* (6–10). Acetylation is another common modification. For example, acetylation of *Vibrio* polysaccharide is required for biofilm formation in *Vibrio cholerae* (11) and acetylation of cellulose is required for surface colonization in *Pseudomonas fluorescens* (12). Acetylation of alginate in *P. aeruginosa* is not only involved in forming the mature biofilm structure (13, 14), but also reduces opsonic killing by phagocytes, reduces susceptibility to enzymatic degradation, and aides in scavenging reactive oxygen species (6, 15–17).

In *P. aeruginosa*, alginate acetylation is known to require four proteins, AlgI, AlgJ, AlgF, and AlgX (18–22). Belonging to the membrane-bound *O*-acetyltransferase (MBOAT) family of proteins (18, 21), AlgI is hypothesized to receive an acetyl group from an unknown donor in the cytoplasm and transfer it to AlgJ in the periplasm (23). Although AlgJ exhibits acetylesterase activity *in vitro*, AlgJ is unable to bind or acetylate poly-mannuronate oligomers *in vitro*, suggesting that it is an intermediary in alginate acetylation (20). AlgX is a periplasmic acetyltransferase that can remove an acetyl group from an artificial donor and transfer it onto mannuronate polymer *in vitro.* In addition, chromosomal mutation of AlgX residues required for its acetyltransferase activity led to production of non-acetylated alginate (19, 20). Thus, it has been hypothesized that a relay takes place to transfer an acetyl group from the cytoplasm to the polysaccharide chain, via AlgI, AlgJ and finally AlgX (18–20). AlgF is also localized to the periplasm and is required for alginate acetylation *in vivo*, but little is known about its structure or function to date (18, 22).

Here, we describe the structure of AlgF determined by CS-Rosetta from chemical shifts data (24) and supported by interproton nuclear Overhauser effect (NOE) analysis. We demonstrate protein-protein interactions *in vitro* between AlgF and AlgJ, as well as AlgF and AlgX, by isothermal titration calorimetry (ITC), and an interaction between AlgF and AlgX identified by co-immunoprecipitation (co-IP) in *P. aeruginosa*. Based on these results, we propose that AlgF functions to mediate interactions between AlgJ and AlgX that are critical for the acetyl relay mechanism required for alginate acetylation, insights that allow us to propose the most detailed model yet of how alginate is modified prior to export.

## RESULTS

### AlgF consists of two β-sandwich domains joined by a short linker region

AlgF is annotated as a periplasmic *O*-acetyltransferase in the *Pseudomonas* Genome Database for several species, including *P. aeruginosa* and *Pseudomonas putida* (25). This annotation is probably due to its demonstrated role in alginate acetylation (18), but sequence alignments with the previously characterized *O*-acetyltransferases AlgJ (20) and AlgX (19, 20) reveal that AlgF lacks similarity to these and other known acetyltransferases. To gain insight into the role of AlgF in alginate acetylation we first determined its structure. Despite exhaustive attempts, the protein proved to be recalcitrant to crystallization, and therefore CS-Rosetta models (26) were generated and validated using NMR spectroscopic techniques.

Uniformly ^1^H,^15^N, and ^13^C labeled *P. aeruginosa* AlgF lacking its signal sequence, AlgF_Pa_ ^30-216^, was expressed and purified. Backbone, triple, and side-chain resonances were assigned by NMR (see Methods) and NOE distance restraints were collected for structural modelling (26–35). AlgF*_Pa_* chemical shifts were assigned to 94.6% completeness. Analysis of the chemical shift data using the chemical shift index (CSI) as calculated by NMRView (28, 30) (Figure S1) indicated that the N- and C-termini were not involved in any stable secondary structural elements as the CSI values were close to zero, as expected for regions with random coil characteristics. Sequence analysis revealed that the protein contains repeating homologous segments, suggesting the presence of a pair of tandem domains joined by a short interdomain region (Figure 1A). The termini and the interdomain region (as well as many loop/turn regions) did not show any long-range NOE assignments (assignments more than four amino acid residues away in sequence) indicating that these regions are only in close contact with their sequential neighbours (Figure S1). Analysis of the folded regions suggested that the structure was all β-strand as reflected by positive CSI scores and a lack of NOE patterns typically found in helical structures (*i.e.* backbone interproton NOEs seen between residues 3 or 4 amino acids apart, i-i+3 and i-i+4 in Figure S1). There is good agreement with the secondary structure observed in the early atomic models calculated using NOE-derived distance restraints (Figure S1, bottom row) and chemical shift data analysis. Despite the quality of the data there were an unacceptable number of clashes and NOE distance restraint violations in ensembles calculated using NOE derived restraints in CYANA (27) (Table 1). This is probably the consequence of the 32% sequence identity between the N- and C-terminal domains (Figure 1A), which results in a high degree of overlap in many chemical shifts resonances. Given the ∼95% completeness of the chemical shift assignments, these data were therefore used in conjunction with the CS-Rosetta server to calculate structural models (26). As is typical for multidomain proteins, when the chemical shift data for the full-length protein was submitted to CS-Rosetta the algorithm failed to converge on a single structure. However, separating the shift data for the two domains provided well-converged structures that matched the NMR restraint-derived structures (Figure 1B and C) with backbone root-mean-square deviations (RMSD) of 1.6 Å and 1.7 Å between the lowest energy CS-Rosetta and NMR-derived N-terminal and C-terminal domain models, respectively. All calculations using NOE-derived distance restraints gave ensembles consisting of two tandem domains (Figure 1). Four amino acids connect the two domains that we could not resolve structurally. Despite extensive searching, no unambiguous interdomain NOE assignments could be found. These results are either because no interdomain NOEs exist, or the structural and sequence similarity between the domains resulted in overlapping chemical shift assignments obscuring the inter-domain NOE assignments.

**Figure 1:**
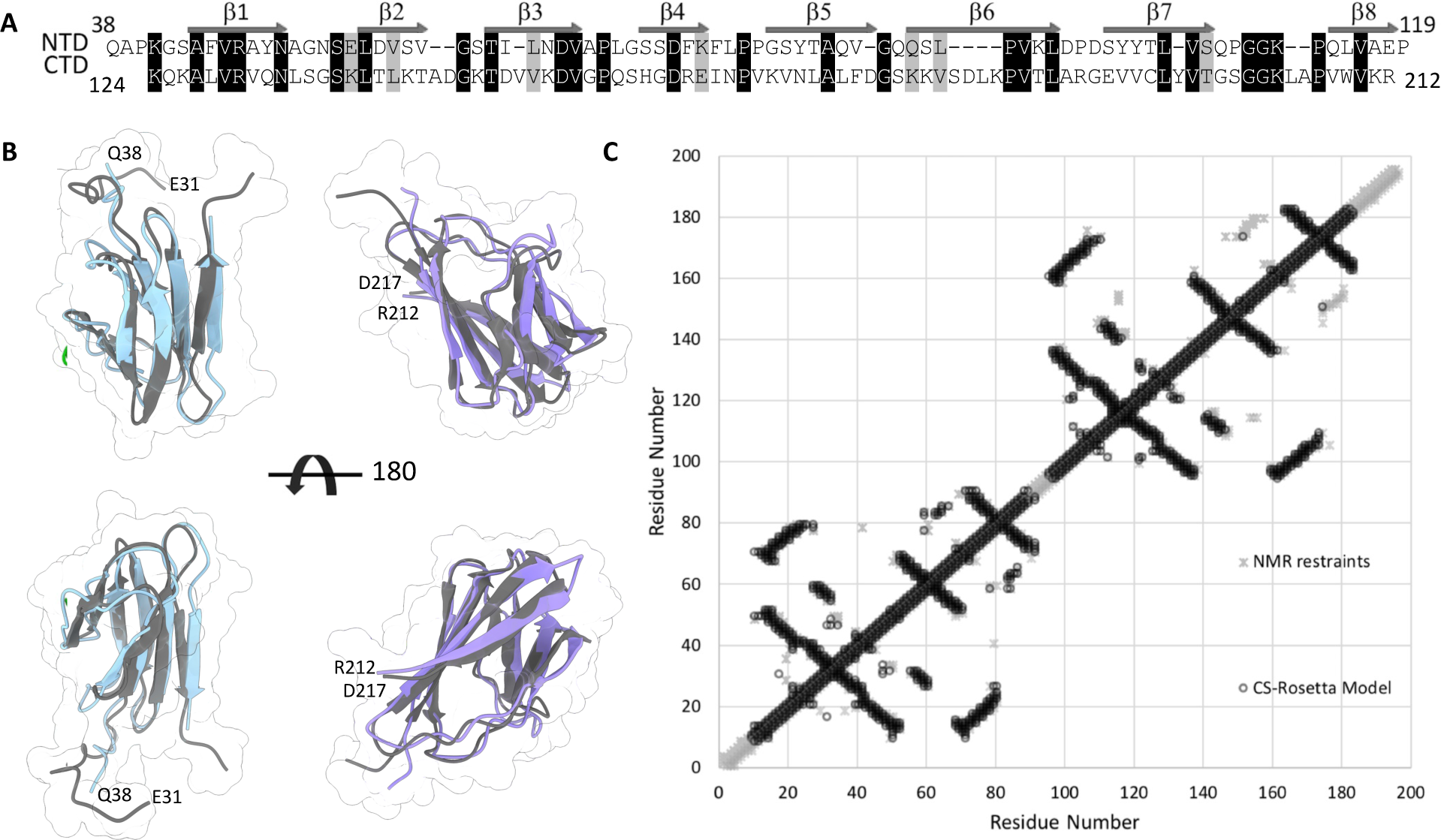
Comparison of CS-Rosetta and NOE derived structures. (A) Sequence alignment of N-terminal (residues 38-119) and C-terminal (residues 124-212) domains of AlgF*_Pa_*^30-216^. Identical residues are shaded in black, similar residues are shaded in grey. Secondary structure elements are shown for each domain with secondary structure elements labeled across the top. (B) Superposition of the lowest energy CS-Rosetta (N-terminal blue, C-terminal purple) and NOE-based CYANA structures (lowest energy model in dark grey). The terminal residues are labeled. (C) Contact map analysis between the CS-Rosetta and NMR determined models. The inter-residue contacts observed in the NMR restraint derived model and the CS-Rosetta derived model are shown in grey asterisks and black circles, respectively. The inter-residue contacts are closely mirrored in structures determined by either method showing the similarity shared by the structures determined by independent methods.

**Table 1:** Summary of model statistics for AlgF ^30-216^.

|  | CS-Rosetta |  | NOE derived |  |
| --- | --- | --- | --- | --- |
|  | N-terminal domain | C-terminal domain |  |  |
| Experimental restraints |  |  |  |  |
| CS-Rosetta input |  |  | CYANA input |  |
| Residue range submitted | 30-119 | 120-222 | Assigned chemical shifts (%) | 2043 (94.6) |
| Folded Core<sup>a</sup> | 38-119 | 124-212 | Short range restraints<sup>b</sup> | 2267 |
| <sup>13</sup>C<sup>a</sup> shifts | 88 | 103 | Medium range restraints<sup>b</sup> | 438 |
| <sup>13</sup>C<sup>b</sup> shifts | 79 | 94 | Long range restraints<sup>b</sup> | 2145 |
| <sup>13</sup>C' shifts | 84 | 100 | Angle restraints | 238 |
| <sup>15</sup>N shifts | 75 | 95 | H-Bond restraints | 130 |
| <sup>1</sup>H<sup>N</sup> shifts | 75 | 95 |  |  |
| <sup>1</sup>H<sup>a</sup> shifts | 98 | 111 |  |  |
| Assigned chemical shifts (%) | 914 (93) | 1148 (94) |  |  |
| Ensemble RMSD values |  |  |  |  |
| Residue range | 44-119 | 127-211 | 42-120<sup>c</sup> | 125-211<sup>c</sup> |
| Backbone atoms Å (# atoms) | 1.38 (304) | 1.59 (340) | 0.32 (316) | 0.24 (348) |
| Heavy atoms Å (# atoms) | 6.23 (1116) | 8.83 (1300) | 10.77 (1136) | 11.67 (1340) |
| Model Quality Measurements |  |  |  |  |
| Ramachandran statistics<sup>d</sup> |  |  |  |  |
| Most favoured regions (%) | 98 | 95 |  | 77.6 |
| Additionally allowed regions (%) | 2 | 4 |  | 21.9 |
| Generously allowed regions (%) | 0 | 0 |  | 0.4 |
| Disallowed regions (%) | 0 | 1 |  | 0.2 |
| Other Quality statistics |  |  |  |  |
| Bad contacts/1,000 residues | 1 | 0 |  | 33 |
| Violated restraints | N/A | N/A | Distance > 0.5 Å<br>Angle > 10° | 91<br>0 |

<sup>a</sup> Residues that were determined by CS-Rosetta to be part of the folded core (regions with chemical shift values close to that of a random coil were truncated by the algorithm prior to fragment generation and model generation)
<sup>b</sup> Definitions for restraint ranges: short range $|i-j| < 1$ , medium range $1 < |i-j| < 5$ , long range $|i-j| > 5$ .
<sup>c</sup> The domains are aligned separately as the lack of inter domain contacts generates highly heterogeneous models with regard to relative domain positioning.
<sup>d</sup> Results for CS-Rosetta models from the PDB validation report and for the CYANA based models from the CYANA output. The statistics for the CYANA model are for the full-length protein.

Both the N- and C-terminal domains of AlgF*_Pa_* form an 8-stranded β-sandwich with backbone ensemble RMSDs for the CS-Rosetta determined models of 1.4 Å and 1.6 Å, respectively (Figures 2A, S2 and Table 1). The AlgF*_Pa_* β-sandwich has two distinct sides; one side is flatter with longer strands (β1, β4, β7, and β8), while the other side has a slight curve with shorter strands and longer loops (β2, β3, β5, and β6) (Figure S2). This asymmetric β-sandwich with one flat face and one curved face was found in both the CS-Rosetta and NOE derived models (Figure 1). The N- and C-terminal domains are structurally similar to each other and superpose with an average ensemble backbone RMSD of 2.1 Å (Figure 2A). Superimposition of the N- and C-terminal domains with the AlphaFold (36) model of AlgF*_Pa_* which predicts the structure with high confidence (Figure S3), reveals a backbone RMSD of 1.5 and 1.6 Å, respectively (Figure 2B). Overall, comparison to the AlphaFold model suggests that the N- and C-terminal domains interact to form a compact structure.

**Figure 2:**
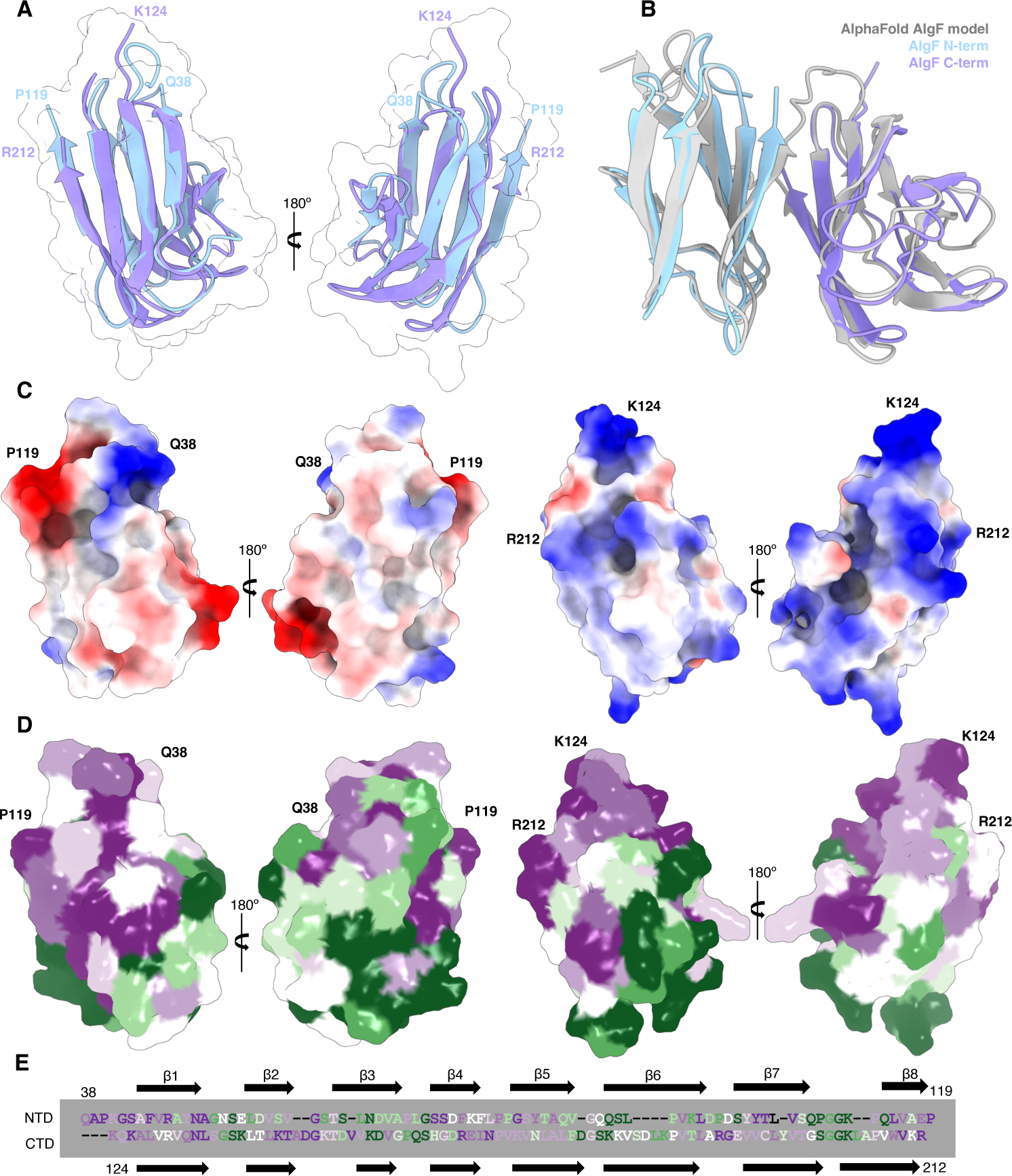
Comparison of the AlgF*_Pa_* N- and C-terminal domains. (A) Superposition of N-(model 9) and C-terminal (model 1) domains with an RMSD of 1.35 Å. The average backbone RMSD for the N- and C-terminal domain ensembles is 2.1 Å. (B) Structural superimposition of the N- and C-terminal domains of AlgF*_Pa_* with the AlphaFold model of AlgF*_Pa_*. (C) Electrostatic potential surface representations of the N- and C-terminal domains (left and right, respectively). Shown in same orientation as panel A. The coulombic potential range from −10 (red) to 10 (blue) kcal/(mol × *e*). (D) ConSurf analysis of conserved residues in the N- and C-terminal domains (left and right, respectively). Shown in same orientation as panels A & C. Surface representation colored by level of conservation (green variable to purple highly conserved). (E) Sequence alignment (as in Figure 1) coloured by conservation level.

To gain insight into the function of AlgF*_Pa_*, the surface characteristics of each domain were further analyzed with respect to charge and sequence conservation. The theoretical pI values were calculated to be 4.60 and 9.61 for the N- and C-terminal domains, respectively, and these differences are reflected in the calculated coulombic surface potential maps (Figure 2C). The N-terminal domain of AlgF*_Pa_* has patches of negatively and positively charged residues, while the C-terminal domain is mostly positively charged. Using the ConSurf server (37), we identified highly conserved surface patches on both N- and C-terminal domains (Figure 2D and 2E). When conservation was analyzed in the context of the compact AlphaFold2 AlgF*_Pa_* structure, highly conserved patches on both domains become buried, further suggesting that the two domains interact (Figure S4). Specifically, the highly conserved residues Arg46, Ala50, Ser72, Ser73, and Val105 on the N-terminal domain and Asn132, Leu133, Val209, and Arg211 on the C-terminal domain are buried in the AlphaFold AlgF*_Pa_* model (Figure S4A). Of these highly conserved residues, Arg46 on the N-terminal domain and Arg211 and Asn132 on the C-terminal domain are involved in electrostatic interactions (Figure S4B). The most electropositive/electronegative regions of AlgF are not involved in mediating interactions between the two domains. The highly conserved residues Val105 on the N-terminal domain and Val209 and Leu133 on the C-terminal domain are involved in hydrophobic interactions (Figure S4C). Less conserved residues, Tyr48 and Val115 on the N-terminal domain and Tyr196 and Val207 on the C-terminal domain, are also involved in hydrophobic interactions between the two domains (Figure S4C). Analysis of the AlgF*_Pa_* AlphaFold model by the Proteins, Interfaces, Surfaces, and Assemblies (PISA) server (38) also predicts that the N- and C-terminal domains interact, with an interaction interface buried surface of 752.4 Å^2^. PISA indicates that the following residues become buried or are solvent inaccessible in the interaction interface: Arg46, Tyr48, Ala50, Ser72, Ser73, Val105 and Val115 on the N-terminal domain are buried, while Leu133, Tyr196, Val207, Val209, and Arg211 on the C-terminal domain are buried (Figure S4D). These data support a compact AlgF structure where the N- and C-terminal domains interact.

### Structurally similar proteins are involved in protein-protein and/or protein-ligand interactions

The AlgF*_Pa_* CS-Rosetta and AlphaFold2 models were submitted individually to the DALI server (39, 40) to identify structurally similar proteins. Based on the nature of ligands bound, we identified four different classes that represent most of the hits identified: 1, protein-binding (e.g. PepT2 solute carrier family 15 extracellular domains, BcpA); 2, cholesterol-binding (e.g. pneumolysin); 3, carbohydrate-binding (e.g. rhamnogalacturonase); and 4, hormone/vitamin-binding (e.g. transthyretin) (Figure 3). Although all four classes contain structurally similar β-sandwich domains, each class is capable of binding chemically distinct ligands.

**Figure 3:**
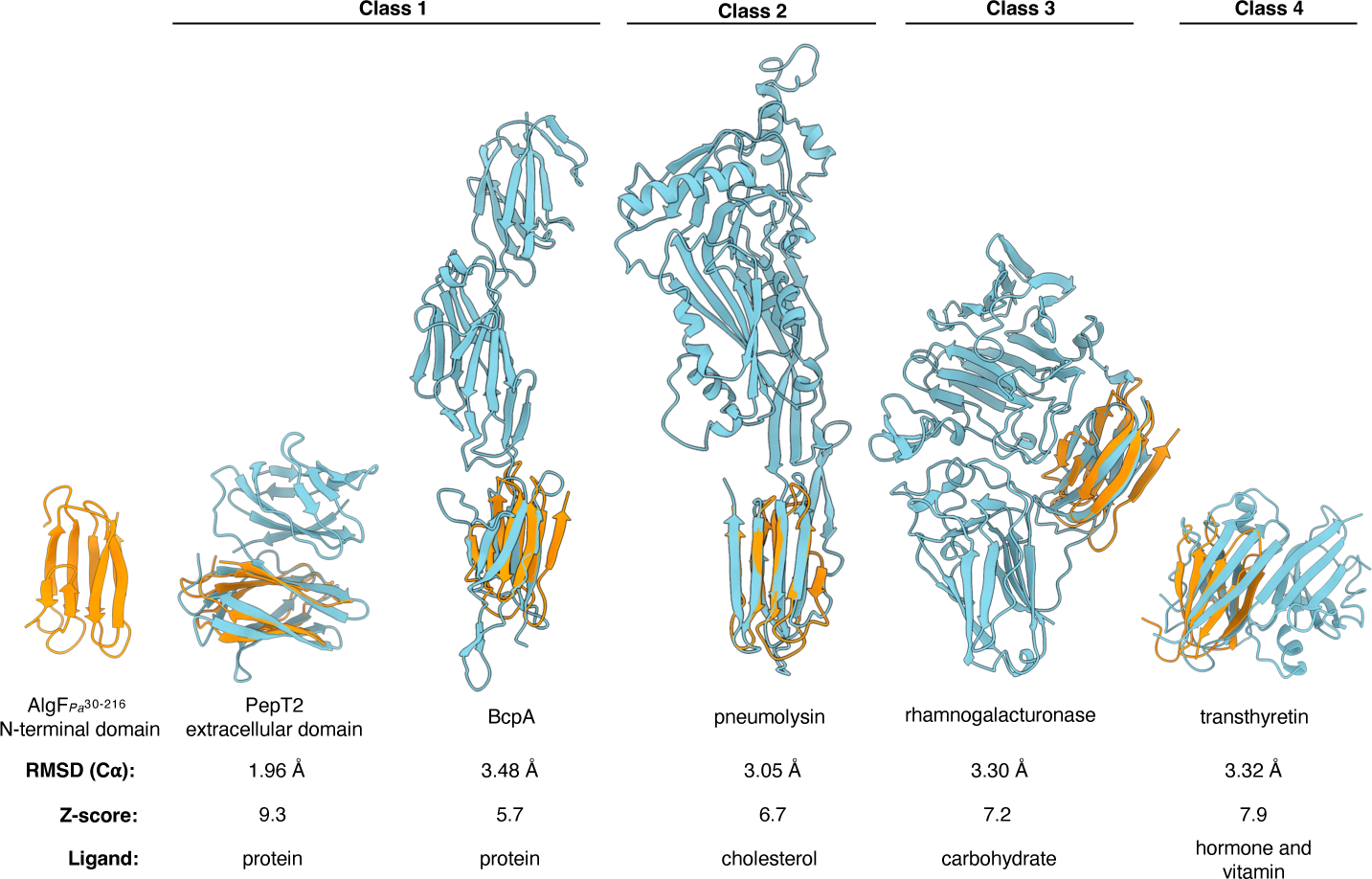
Structural homologs of AlgF bind a variety of ligands. Structural superimposition of AlgF*_Pa_* N-terminal domain (orange) with protein structures (teal) representing most (> ∼60%) of the overall hits identified using the DALI server to the N-terminal domain of AlgF*_Pa_*. C_α_ atom RMSD values and Z-scores for homologous structures are as indicated. The ligand or proposed ligand of the β-sandwich domain for each protein hit is indicated. Classes were defined by the nature of the ligand. The same hits were found using DALI when the C-terminal domain of AlgF*_Pa_* and the AlphaFold model of AlgF*_Pa_* were used as the search structure. The DALI Z-scores represents a similarity score that considers the distances between C_α_ atoms between two proteins. (PDB codes left to right: 5A9H, 3KPT, 5AOE, 2XHN, 6R66).

Class 1 proteins have the most structural homology to AlgF*_Pa_*. Within class 1, solute carrier family 15 proteins (PepT1 and PepT2 (41)) are involved in oligopeptide transport across mammalian cell membranes and the extracellular domains of these proteins bind trypsin and recruit it to the site of dietary peptide uptake. PepT1 and PepT2 extracellular regions consist of two tandem β-sandwich domains joined by a short linker, and most closely resemble the β-sandwich domain structure of AlgF*_Pa_* (Figure S5A). However, only PepT2’s domains appear to interact to form a compact particle, similar to AlgF*_Pa_* (Figure S5A). Despite the similar structures, PepT1 and PepT2 share 19% and 14% amino acid sequence identity, respectively, to the AlgF N-terminal domain. Sequence comparisons to the AlgF*_Pa_* C-terminal domain are similar; 14% and 16% for PepT1 and PepT2, respectively. The bacterial pilin adhesion protein BcpA is critical for maintaining cell-cell contacts through intramolecular amide bonds, resulting in pili fibre formation on the cell surface (42). While BcpA has three tandem β-sandwich domains which are structurally similar to the N-terminal and C-terminal domains of AlgF*_Pa_*, the amino acid sequence identity between any two of the aligned domains is less than 6%. Class 2 consists of cholesterol-binding proteins including pneumolysin, a toxin found in pathogenic *Streptococcus pneumoniae* that forms a pore in eukaryotic membranes (43). Only the cholesterol-binding domain of the toxin forms a β-sandwich that is structurally similar to the N-terminal and C-terminal domains of AlgF*_Pa_*. Domain II of rhamnogalacturonase in class 3 is suggested to be involved in oligosaccharide binding and resembles a single domain of AlgF (22, 23). The class 4 protein transthyretin is named for its role in transporting thyroxine and retinol in serum (44). While transthyretin consists of β-sandwich structures, the domains are oriented differently compared to AlgF*_Pa_* (Figure S5B) (44). Most of the remaining unclassified hits (*i.e.* not belonging to classes 1-4) also have a role in binding small molecules and have little similarity to the structure of AlgF. In the case of multi-domain proteins identified by the DALI search, the AlgF-like β-sandwich domain functions to bind small molecules, while the rest of the respective protein carries out the biological function. Since AlgF lacks any ancillary domains, we hypothesize that it most likely functions solely to bind either a small molecule or protein ligand.

### In vitro binding analyses reveal direct interactions between AlgF, AlgJ, and AlgX

Since class 1 proteins with a β-sandwich structure are involved in binding proteins and AlgF has been proposed to interact with the acetylation machinery (45), we next investigated whether AlgF is involved in protein-protein interactions. Binary interactions between *P. putida* homolog constructs AlgF_Pp_ ^30-215^, AlgJ ^75-370^, and AlgX_Pp_ were probed *in vitro* using isothermal titration calorimetry (ITC). These proteins could be obtained in higher yields and were more stable than their *P. aeruginosa* counterparts thus enabling the ITC experiments (Figure 4). We found that AlgJ*_Pp_* and AlgX*_Pp_* bind to AlgF*_Pp_* with K_d_ values of 86 ± 12 μM and 178 ± 5 μM, respectively. Titration of AlgX*_Pp_* into AlgJ*_Pp_* demonstrated a decreasing heat of enthalpy as the titration proceeded, however the data could not be reliably fit as the heats of emission were small and the titration did not reach saturation. The data suggest a weak interaction with a dissociation constant in the mM range.

**Figure 4:**
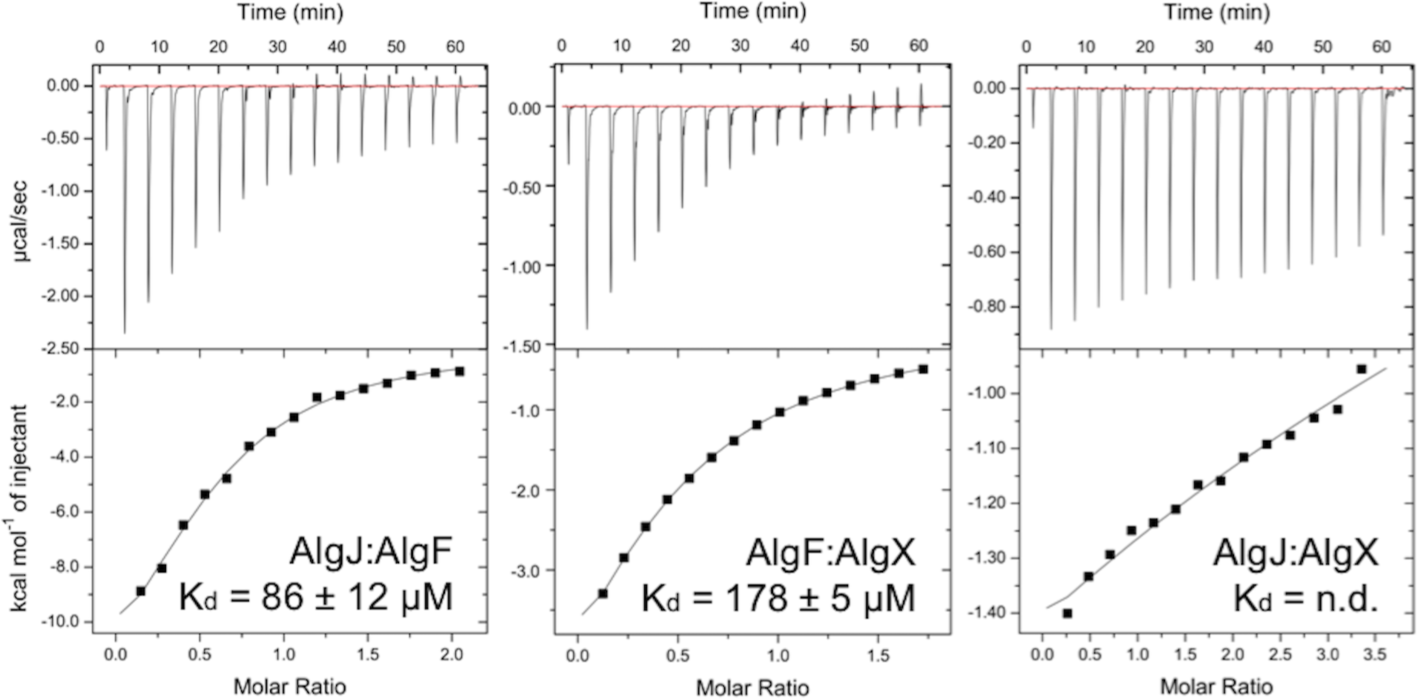
ITC analyses of AlgF*_Pp_*^29-215^ protein-protein interactions with AlgJ*_Pp_*^75-370^ and AlgX*_Pp_*. Protein-protein interactions were investigated by binary titrations of AlgJ:AlgF (left), AlgF:AlgX (centre) and AlgJ:AlgX (right) using ITC. Data were fit and analyzed with the Origin software package as shown, and K_d_ values were determined. The AlgJ:AlgX interaction (right) was too weak to be analyzed.

### AlgF is not an acetyltransferase and does not bind alginate

In Δ*algF* strains of *P. aeruginosa*, previous studies found that alginate exopolysaccharide is produced but not acetylated (18). The fold of AlgF did not suggest a particular enzymatic role and the 3D structure alone was not sufficient to provide insight into the function of AlgF. Unlike AlgJ and AlgX, AlgF shared no structural similarities with acetyltransferase enzymes. As AlgF is required for alginate acetylation, we investigated if this requirement was the result of a previously uncharacterized enzymatic activity.

An assay was carried out to determine whether AlgF*_Pp_* had acetylesterase activity, the first step in the acetyltransferase reaction. Removal of an acetate group from the pseudosubstrate 3-carboxyumbelliferyl acetate results in release of a fluorescent product, and reaction progress can be monitored through fluorescence spectroscopy. The acetylesterase activity of *P. aeruginosa* AlgF*_Pp_*, AlgX*_Pp_*, and AlgJ*_Pp_* was measured independently, in combination, and in the presence of an acetyl group acceptor (a non-acetylated mannuronic acid decasaccharide; ManA_10_). No acetylesterase activity was observed for AlgF*_Pp_* (Figure 5). Addition of AlgF*_Pp_* to either AlgJ*_Pp_* or AlgX*_Pp_* did not increase observed acetylesterase activity, with or without presence of ManA_10_ (Figure 5). Similarly, addition of AlgF*_Pp_* to a combination of AlgJ*_Pp_* and AlgX*_Pp_* did not result in an increase acetyltransferase activity, with or without presence of ManA_10_, suggesting that the presence of AlgF does not influence overall acetylesterase activity. Therefore, the proposed formation of an acetylation machinery complex, although required for alginate acetylation *in vivo*, does not affect the acetylesterase activity of AlgJ or AlgX *in vitro*.

**Figure 5:**
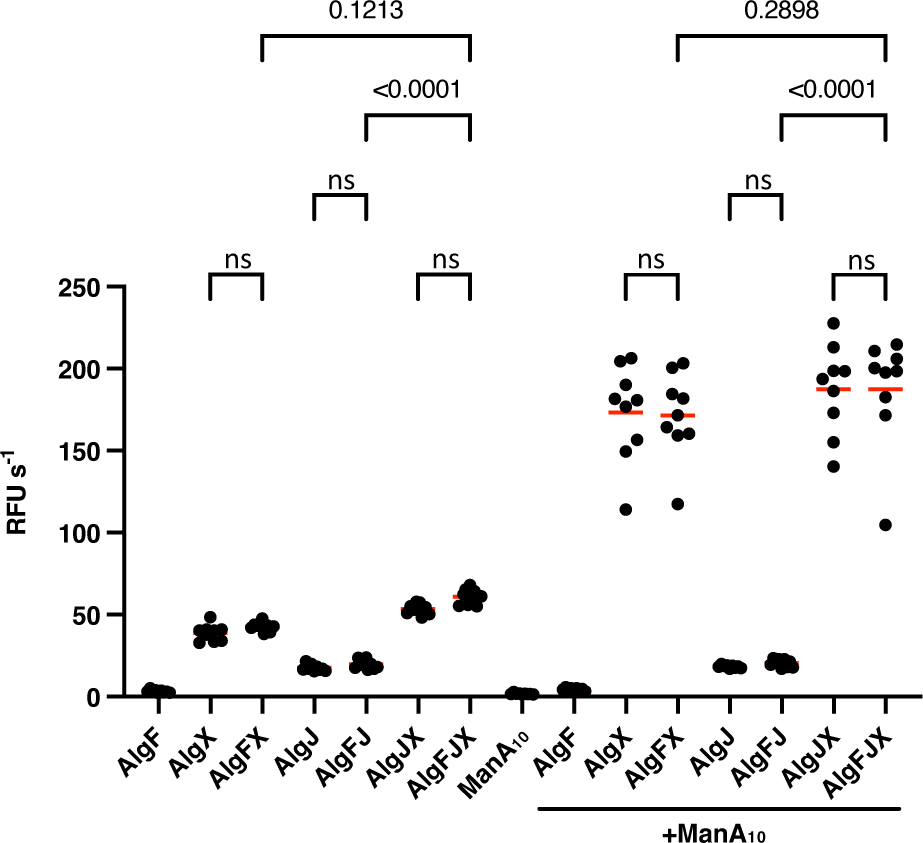
Acetylesterase activity of AlgF and the putative acetylation complex. Enzyme activity was measured by hydrolysis of acetate from the pseudosubstrate ACC. Release of the 7-hydroxycoumarin-3-carboxylic acid fluorescent product was measured (λ_ex_ = 386 nm and λ_em_ = 446 nm). Values represent three technical replicates for three biological replicates. Red lines represent the mean. The reactions were performed with the addition of 2 mM ACC to 5 μM of each protein. Buffer contained 50 mM sodium HEPES pH 7.6 and 75 mM NaCl at 25 ℃. ManA_10_ denotes the addition of 1 mg/mL chemically synthesized polymannuronate decasaccharide.

Even though AlgF*_Pp_* demonstrated a lack of acetylesterase activity, some of the AlgF-related ß-sandwich proteins bind sugars/small molecules. Thus, the ability of AlgF*_Pa_* to bind carbohydrate polymer was investigated using an electrospray ionization mass spectrometry (ESI-MS) binding assay. Nine mannuronic acid oligomers ranging from 4 to 12 sugars in length (ManA_4_ to ManA_12_) were tested. AlgF*_Pa_* did not display any affinity for the oligomers tested. The approximate K_a_ values measured were less than 500 M^-1^ for all oligomers tested (Table S1). This is similar to what was observed previously for AlgJ (20). In contrast, AlgX binds to mannuronic acid oligomers in a length dependent manner (20). As the data suggest that AlgF does not bind alginate in isolation, it most likely functions as a protein-protein interaction mediator between AlgJ and AlgX.

### AlgIJFX form a complex in P. aeruginosa

The *in vitro* binding data obtained by ITC suggests that AlgF interacts with AlgX and AlgJ, and that AlgF may mediate the formation of a complex when all three proteins are present. To examine whether these findings also hold true in *P. aeruginosa*, we performed co-immunoprecipitation (co-IP) experiments from lysed *P. aeruginosa* cells expressing vesicular stomatitis virus glycoprotein (VSV-G) tagged AlgX, AlgJ, or AlgF. The VSV-G sequence was introduced at the C-terminus of each gene directly on the chromosome of PAO1 Δ*wspF* P_BAD_*alg* (46). In this strain, the native *algD* promoter has been replaced by *araC*-P_BAD_, allowing for high level inducible expression of the *algD* operon in the presence of arabinose. Clarified lysates were applied to agarose resin conjugated to anti-VSV-G monoclonal antibodies, and the elution from the resin after washing was analyzed by Western blot using protein-specific polyclonal antibodies. The corresponding untagged protein was used as a negative binding control. When AlgX_Pa_ ^C-VSV-G^ was supplied as the bait, AlgF*_Pa_* was identified as an interaction partner (Figure 6A). AlgF*_Pa_* was not identified in the elution from the negative control, indicating that the observed interaction is not due to non-specific binding with the co-IP resin (Figure 6A). To confirm this finding, co-IP eluates from six independent co-IP experiments using AlgX_Pa_ ^C-VSV-G^ as the bait were analyzed by ESI-MS. AlgF_Pa_, as well as the previously identified interaction partner AlgK*_Pa_* (47, 48), were significantly enriched in the AlgX*_Pa_*^C-VSV-G^ eluate versus the untagged negative control (Figure 6B), confirming the interaction between AlgX*_Pa_* and AlgF*_Pa_*. When AlgF*_Pa_*^C-VSV-G^ was supplied as the bait, no co-eluting proteins were identified suggesting that addition of the VSV-G tag to AlgF*_Pa_* may have disrupted the stability or function of AlgF*_Pa_*.

**Figure 6:**
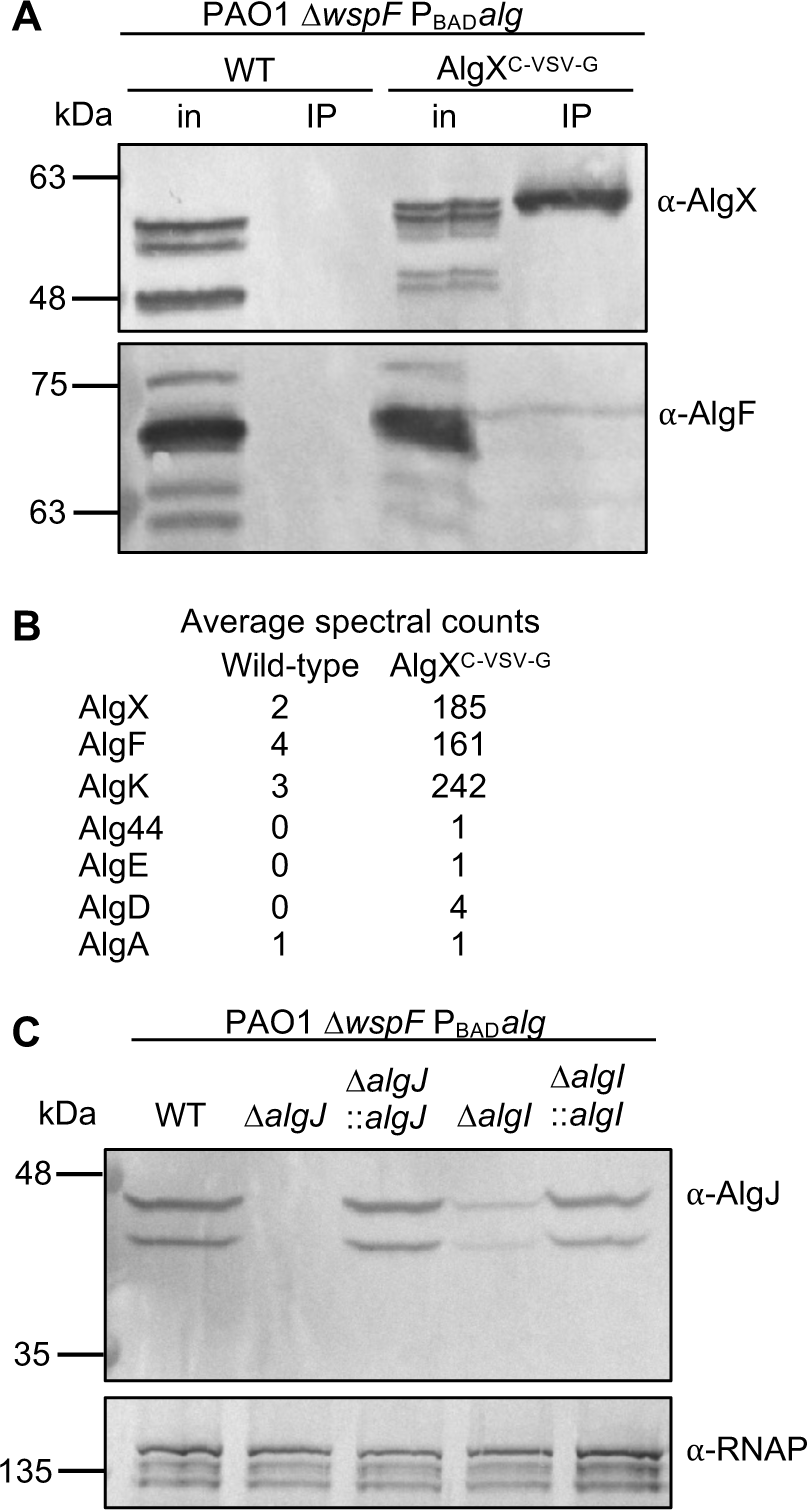
AlgX-AlgF and AlgI-AlgJ interact in *P. aeruginosa*. (A) Co-IP from whole cell lysates of *P. aeruginosa* expressing VSV-G-tagged AlgX as the bait. Proteins applied to the anti-VSV-G co-IP resin (input, in) and the elution from the resin after washing (IP) were analyzed by Western blot using AlgX- and AlgF-specific antibodies. A strain expressing untagged AlgX was used as a negative binding control. (B) Analysis of co-IP eluates from the experiment described in panel *A* by ESI-MS. Spectral counts were the average of six independent co-IP experiments using AlgX^C-VSV-G^ as the bait. Only alginate biosynthetic proteins identified with a minimum of one spectral count in this analysis are listed. (C) Western blot analysis of whole cell lysates of the indicated *P. aeruginosa* strains using AlgJ-specific antibodies. Antisera recognizing the β-subunit of RNA polymerase was used as a loading control.

Co-IP could not be performed successfully with AlgI*_Pa_* as the bait due to the instability and aggregation of AlgI*_Pa_* after solubilization from *P. aeruginosa* membranes. Attempts to optimize extraction using various detergents were unsuccessful, precluding the ability to generate an AlgI*_Pa_*-specific polyclonal antibody. While we were able to generate a VSV-G-tagged construct of AlgI*_Pa_* that could complement acetylation in an *algI* deletion mutant, AlgI*_Pa_* also aggregates in Laemmli buffer under all conditions tested, preventing its detection by Western blot. To determine whether the destabilization of AlgI*_Pa_* during the detergent extraction step of co-IP experiments would have an effect on the stability of other acetylation proteins, a mutual stability analysis was performed using a Δ*algI* variant to mimic the loss of AlgI*_Pa_* due to detergent extraction-mediated aggregation. When expression of AlgJ*_Pa_* was analyzed by Western blot, a significant reduction in steady-state protein levels was observed in the Δ*algI* background versus wild-type (Figure 6C). Complementation of *algI* at the neutral *attTn7* site on the *P. aeruginosa* chromosome restored AlgJ*_Pa_* to wild-type levels (Figure 6C), suggesting that the reduction in whole-cell AlgJ*_Pa_* levels was due specifically to the deletion of *algI*. Indeed, when co-IP experiments were performed with AlgJ_Pa_ ^C-VSV-G^ as the bait, no interaction partners were identified, likely due to AlgI*_Pa_* aggregation after solubilization of the membranes and the resultant effects on the stability of AlgJ*_Pa_*. Overall, these data support the presence of a physiological interaction between AlgX*_Pa_* and AlgF*_Pa_* and imply also that AlgJ*_Pa_* and AlgI*_Pa_* interact based on the observed stability requirement of AlgJ*_Pa_* for AlgI*_Pa_*.

### Predictive modelling of the AlgKXF and AlgIJ complexes

The structure of the AlgKX*_Pp_* complex has been previously determined experimentally (PDB: 7ULA) and AlphaFold2 was shown to accurately predict the structure/interaction interface of the AlgKX*_Pp_* complex across different *Pseudomonas* species (48). Building on the success of the AlphaFold2 model of AlgKX*_Pp_*, we next sought to determine whether the program could predict how AlgF*_Pp_* may interact with AlgX*_Pp_* and AlgK*_Pp_.* Using this approach, we were able to generate a high-confidence model of the AlgKXF*_Pp_* complex involved in alginate modification and export (Figures 7A and S6). The predicted AlgKXF*_Pp_* complex shows that AlgK*_Pp_* and AlgX*_Pp_* maintain the same interaction interface as previously reported in the experimentally determined structure (48). Analysis of the AlgKXF*_Pp_* model by the PISA server (38) suggests that AlgF*_Pp_* only interacts with AlgX*_Pp_*. The modelled AlgXF*_Pp_* interaction interface was calculated to have a buried surface area of 1545 Å^2^ mediated by 12 hydrogen bonds and six salt bridges (Figure 7B). Most notably, ten out of eleven predicted interacting residues on AlgF*_Pp_* interact with AlgX*_Pp_* using their side chain atoms and all eleven residues are present on coil regions of AlgF*_Pp_*. Three AlgF*_Pp_* interaction interface residues are present on the N-terminal domain (Asp29 and Tyr33 involved in hydrogen bonding and Lys39 involved in both hydrogen bonding and salt bridge interactions), while the remaining nine are present on the C-terminal domain (Gln123, Lys124, Asn163, Pro164, Lys166, Glu187, Arg187 and Glu190) (Figure 7B). Thus, both the N- and C-terminal domains are predicted to be required for the interaction with AlgX*_Pp_*. In AlgF*_Pa_*, these interaction interface residues correspond to Asp29, Tyr34, Lys40, Gln124, Lys125, Asn164, Pro165, Lys167, Ala188, Arg189, and Glu191. Of the nine residues that are present in the AlgF*_Pa_* experimentally determined structure, four of these residues are highly conserved, four are conserved, and one is less conserved (Figure 2E). Building on this model, we next attempted to model the AlgKXFJ*_Pp_* complex using AlphaFold2. To expand this prediction to include AlgJ*_Pp_*, due to the 1400 residues limitation in AlphaFold2, we only included the C-terminal region of AlgK*_Pp_* that binds to AlgX*_Pp_* when predicting the AlgKXFJ*_Pp_* complex. Although the predicted models revealed a consistent AlgKXF*_Pp_* interaction interface (as shown in Figure 7), the predicted alignment error (PAE) plot revealed the interaction of AlgJ*_Pp_* with AlgKFX*_Pp_* could not be accurately predicted as the PAE values for AlgJ*_Pp_* were estimated to be >25Å (Figure S7). Thus, we were unable to generate a high-confidence model to illustrate how AlgF*_Pp_* and AlgJ*_Pp_* may interact.

**Figure 7:**
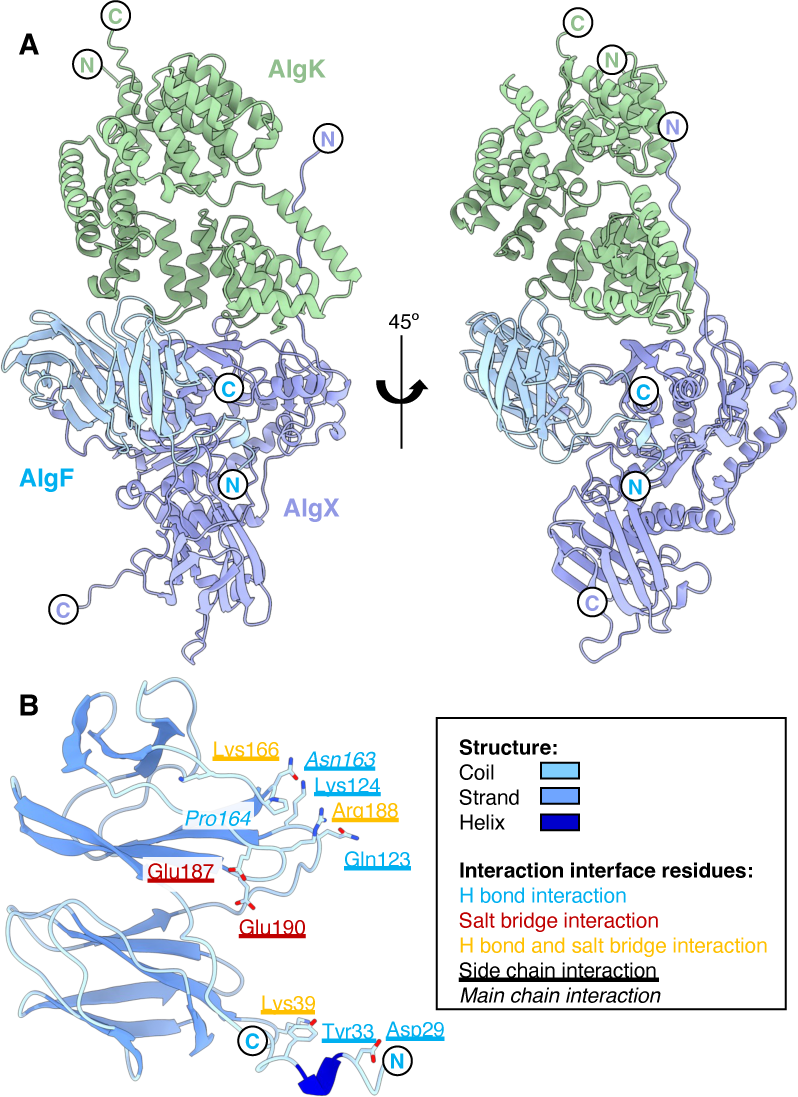
Model of the AlgKXF*_Pp_* complex involved in alginate acetylation and export. (A) AlphaFold2 model of the *P. putida* AlgKXF complex (AlgK*_Pp_*, green; AlgX*_Pp_*, periwinkle; AlgF*_Pp_*, light blue). (B) AlgF*_Pp_* residues involved in the interaction interface with AlgX*_Pp_*. The model of AlgF*_Pp_* is coloured by secondary structure (coil, light blue; strand; blue; helix, dark blue). Residues involved in hydrogen bonding interactions, salt bridge interactions, or hydrogen bonding and salt bridge interactions are represented by blue, red, and yellow text, respectively. Residues that are underlined represent side chain interactions and residues that are italicized represent main chain interactions. In both panels the N - and C-termini are represented by N and C, respectively.

Given that our mutual stability data from *P. aeruginosa* suggests an interaction between AlgI*_Pa_* and AlgJ*_Pa_,* we also used AlphaFold to model the AlgIJ*_Pp_* complex. A high-confidence model of the AlgIJ*_Pp_* complex was generated for this part of the predicted acetylation complex (Figure S8). This complex suggests that it is predominantly the transmembrane domain of AlgJ that interacts with AlgI. Analysis by PISA reveals that the interaction interface area between AlgI and AlgJ is 2122 Å^2^, mediated by ten hydrogen bonds and three salt bridges. The N-terminal helix of AlgJ is inserted into the inner membrane and is predicted to pack between the first and last helices of AlgI (Figure S8).

## DISCUSSION

In this study, we present the structure of AlgF and, using *in vitro* studies coupled with analyses from *P. aeruginosa*, establish that the proteins involved in alginate acetylation (AlgI, J, F, and X) interact to form an acetylation complex that is linked to the outer membrane export machinery *via* an interaction between AlgX and AlgK. We found that AlgF consists of two β-sandwich domains joined by a linker, and our functional characterizations suggest that AlgF is unlikely to function as an alginate acetyltransferase as it lacks acetylesterase activity and is unable to bind alginate *in vitro*. Considering that most of its structural neighbours are involved in protein-ligand interactions, we propose that AlgF functions as a protein-protein interaction mediator within the alginate biosynthetic system to coordinate an AlgJFX periplasmic acetylation complex.

A search for AlgF structural homologs using the DALI server revealed proteins involved in ligand binding. The tandem two-domain architecture of AlgF resembles the binding module of the Eukaryotic peptide transporter PepT2, which mediates an interaction between trypsin and the membrane transporter domain (41). This functionality could be mirrored with AlgF localizing and/or strengthening interactions between the acetyltransferase proteins and the rest of the alginate biosynthetic complex. As evidence of this role for AlgF, we have been able to establish that AlgF binds to both AlgX and AlgJ *in vitro*. We originally hypothesized that one domain of AlgF binds AlgJ at the inner membrane while the other binds to AlgX however, structural prediction of the AlgKXF*_Pa_* model suggests that both domains are required for its interaction with AlgX. Thus, further binding studies in conjugation with targeted mutagenesis of AlgF are required to ascertain which regions of AlgF are involved in interacting with either AlgX or AlgJ. Previous studies have demonstrated that alginate is produced but not acetylated when either AlgJ or AlgX enzymatic activity is compromised (19, 20). Taking into consideration all the data presented on AlgJFX thus far, this suggests that transfer of an acetyl group from AlgJ to AlgX occurs and is required for polymer acetylation. As no direct interaction between AlgJ and AlgX was observed in this study, we hypothesize that AlgF is necessary to bring AlgJ and AlgX together in close enough proximity for acetyl relay and that lack of AlgF would decouple the acetyltransferase process. AlgI has also been found to be critical for acetylation of the nascent polymer (18, 21, 23). Our mutual stability studies demonstrate that AlgI may be part of the acetylation complex. This hypothesis is further strengthened by our AlphaFold2 model that suggests that AlgI is linked to the acetylation machinery primarily through the transmembrane domain of AlgJ (45). While we have been unable to model the entire AlgIJFX complex, we propose that AlgIJFX serve as an acetyl relay to transfer an acetate from AlgI, across the inner membrane first to AlgJ, then AlgX and finally to the polymer (Figure 8). Comparably, acetylation of cellulose in *P. fluorescens* requires the proteins WssF/H/I/G which are homologous to the alginate proteins, AlgX/I/J/F (49). The AlgF-like protein, WssG, in acetylated cellulose biosynthesis is poorly characterized and its role remains unknown (49). Future experiments along the same lines as those presented here could help establish whether WssG is also a protein-protein interaction mediator and critical for the formation of a WssH/I/G/F complex.

**Figure 8:**
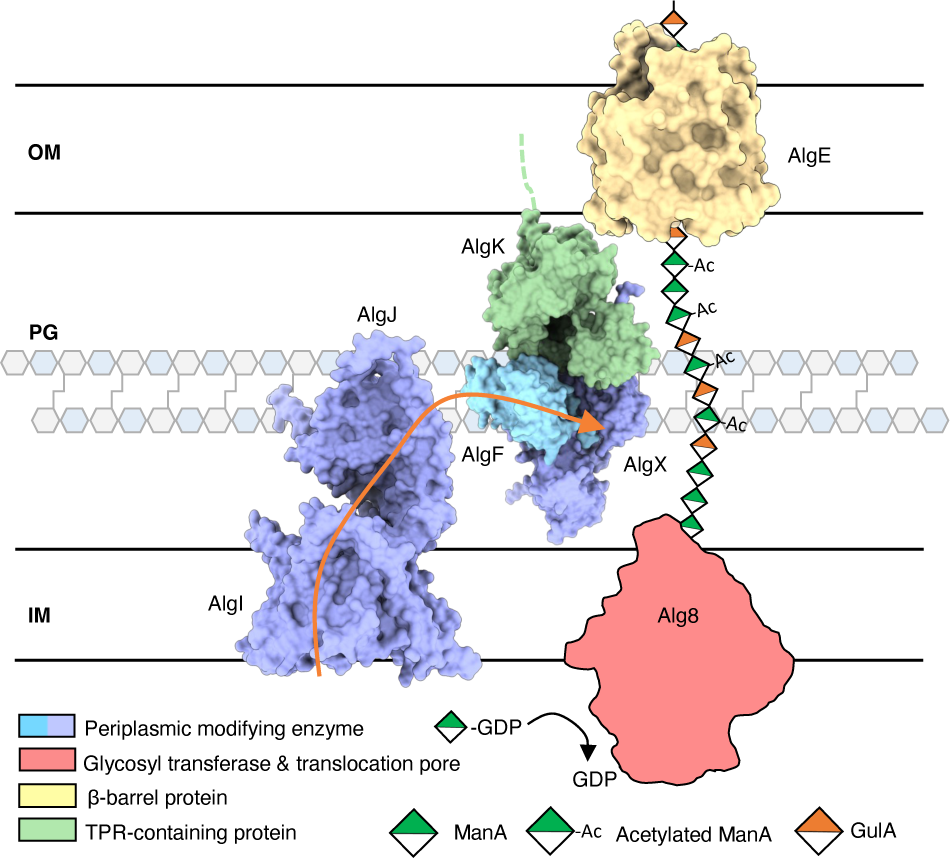
Model of alginate *O*-acetylation machinery. AlgI receives an acetyl group from an as-yet unknown donor in the cytoplasm and the acetate group is passed through the inner membrane and transferred to AlgJ. AlgF mediates interactions in the periplasm allowing for the acetate to be passed from AlgJ to AlgX, where AlgX *O*-acetylates the newly polymerized mannuronic acid. OM, PG, IM denote the outer membrane, peptidoglycan, and inner membrane, respectively. Orange arrow indicates how the acetyl group is transferred between AlgI, AlgJ, and AlgX.

In addition to the AlgI/J/F/X interactions outlined here, previous studies have demonstrated that AlgX and AlgK form a robust complex that couples alginate acetylation and export (48). Furthermore, given that AlgE localization is dependent on the presence of AlgK (50), formation of an AlgEKX outer membrane secretion complex has also been proposed (48). Within the inner membrane, it has been suggested that Alg8 and Alg44 form a synthase co-polymerase complex (51). Alg44’s periplasmic domain has been proposed to interact with AlgX and AlgK (52). These previously established interaction networks highlight the mechanisms required for alginate synthesis. Our ability to observe the well-established AlgX-AlgK interaction using co-IPs reinforces the results obtained here that link AlgX to AlgF and the rest of the acetylation machinery. The interactions between the periplasmic acetylation proteins have been found to be relatively weak by ITC with μM affinities. Thus, it may be possible that this weak binding is an artifact of *in vitro* protein studies of binary rather than native multi-protein interactions. The acetylation proteins may interact with a higher affinity once the rest of the biosynthetic complex and alginate substrate is present, allowing for a more stable complex *in vivo*. This hypothesis is not without precedent in large protein complexes (53–55). Specifically, the VirB type IV secretion system complex functions as a minimal set of VirB7 to VirB10 proteins, while the addition of VirB1 to VirB4 increases the activity of the transport complex dramatically, as well as increases the abundance of macrocomplex protein interactions (54). Weak μM interactions between proteins may also indicate the presence of a transient overall biosynthetic complex (55), where a number of subcomplexes may be present in the alginate biosynthesis pathway with variable intra- and inter-subcomplex affinities. Neither high affinity interactions nor stable complex formation may be needed (or desirable) in order to fine-tune alginate modification, as the degree of acetylation is seen to vary depending on bacterial strain and species or on growth conditions and may even vary over the course of biofilm development, maintenance, and dispersal (56–59). If this is the case, then structural determination of subcomplexes with the alginate biosynthetic system may be more reasonable and realistic compared to determination of the complex in its entirety.

The structural determination of AlgF, structural homology searches, and characterization of protein function have enabled us to identify that AlgF mediates protein-protein interactions in the alginate acetylation machinery. The formation of an AlgIJFX acetylation complex supports the previously proposed relay mechanism for alginate modification where AlgF mediates interactions between AlgJ and AlgX for acetylation of the polymer (18–20). For the first time, we are able to build upon established protein networks described above and show links from the inner membrane complex AlgI/J to AlgX/K at the outer membrane. These complexes are linked via AlgF and join the modification subcomplex in the inner membrane/periplasm with the outer membrane export subcomplex. These data are suggestive of a large protein interaction network and support the hypothesis of a trans-envelope macrocomplex encompassing all of the exopolysaccharide biosynthetic proteins (18, 60). Furthermore, recent studies on the homologous cellulose biosynthetic machinery have shown the presence of an inner membrane subcomplex formed by cytoplasmic, transmembrane, and periplasmic proteins (61–64). Future work with alginate proteins may uncover similar interactions with cytoplasmic components including sugar precursor enzymes, thus expanding our knowledge of how alginate polymerization and precursor biosynthesis may be coupled. Understanding the biosynthesis of exopolysaccharides, including the alginate system, will require further studies of the protein interaction networks not only at a protein-protein level but also at a cellular level. The results presented here provide the first insight into a membrane-spanning polysaccharide secretion complex with significant etiological consequences for patient outcomes in cystic fibrosis.

## METHODS

### Bacterial strains, plasmids, and growth conditions

A detailed list of the bacterial strains and plasmids used in this study can be found in Table S2. All *P. aeruginosa* mutant and complemented strains were derived from PAO1 (65) and were constructed using allelic exchange and miniTn7 mutagenesis (66, 67), as described below. Unless otherwise stated, lysogeny broth (LB) was used for growth of all strains. LB contained, per litre of ultrapure water, 10 g tryptone, 5 g NaCl, and 10 g yeast extract. Vogel-Bonner minimal medium (VBMM) was prepared as a 10× concentrate containing, per litre of ultrapure water, 2 g MgSO_4_·7H_2_O, 20 g citric acid, 100 g K_2_HPO_4_, and 35 g NH_4_HPO_4_, and was diluted to 1× as needed. To prepare solid media, 1.5% (w/v) agar were added to LB or VBMM. Where appropriate, antibiotics were added to growth media. For *E. coli*, 10 μg/ml gentamicin, 100 μg/ml carbenicillin, or 50 μg/ml kanamycin was used. For *P. aeruginosa*, 30 or 60 μg/ml gentamicin was used depending on the application, as described below.

### P. aeruginosa and P. putida gene expression in E. coli

The nucleotide sequences of AlgF from *P. aeruginosa* PAO1 and *P. putida* KT2440, and of AlgX from *P. putida* KT2440 were obtained from the *Pseudomonas* Genome Database (25). AlgF_Pa_ ^30-216^ and AlgF_Pp_ ^29-215^ were PCR amplified from genomic DNA. Primer sequences are as indicated in Table S3. Primers account for entire full-length proteins without the respective N-terminal signal sequence as predicted by SignalP (68) and introduce *Nde*I and *Xho*I restriction sites. The codon-optimized gene for expression in *E. coli* of full-length AlgX*_Pp_* included flanking *Nde*I and *Xho*I restriction sites at the 5’ and 3’ ends, respectively, was synthesized by BioBasic. AlgF_Pa_ ^30-216^ and codon-optimized AlgX_Pp_ were incorporated into the pET24b vector for C-terminal His_6_-tag protein expression. AlgF_Pp_ ^29-215^ was incorporated into the pET28a vector with a 3’ stop codon for N-terminal His_6_-tag protein expression. For each protein construct, *E. coli* BL21 CodonPlus (λDE3) cells (Stratagene) were transformed with expression vector and grown in LB broth containing 50 μg/ml kanamycin at 37 °C. Once the OD600 of the culture reached 0.6, protein expression was induced by the addition of IPTG to a final concentration of 1 mM. The cell culture was incubated at 18 °C for 16 h prior to being harvested by centrifugation at 6700 × *g* for 25 min at 4 °C. Cell pellets were stored until needed at −20 °C. AlgJ_Pa_ ^79-379^ and AlgJ_Pp_ ^75-370^ were expressed as described previously (20).

### Purification of His_6_-tagged protein

The cell pellet from 1 L of bacterial culture was thawed and resuspended in 50 mL of lysis buffer (50 mM Tris pH 8.0, 500 mM NaCl, 0.5 M EDTA, 1 mM DTT, 1 mM PMSF, 2% (v/v) glycerol, 1 mg/mL lysozyme) with one Roche Complete protease-inhibitor cocktail (EDTA-free) tablet. The resuspended pellet was incubated at 4 °C for 30 min. Cells were homogenized at 15000 psi using an Emulsiflex C3 (Avestin Inc.) for 3 passes or until fully lysed. Cell lysate was centrifuged at 20100 × *g* for 20 min at 4 °C to remove cellular debris. The resultant lysate supernatant was loaded onto Ni-NTA resin and washed with 30 column volumes of 50 mM Tris pH 8.0, 500 mM NaCl, 2% (v/v) glycerol, and 30 mM imidazole. Protein was eluted using similar buffer with 300 mM imidazole and concentrated by centrifugation with 4 kDa (AlgF) or 30 kDa (AlgJ and AlgX) cutoff Vivaspin Turbo centrifugal concentrators (Sartorius). His_6_-tagged protein was further purified using a HiLoad 16/60 Superdex 200 prep-grade size exclusion column (GE Healthcare) in 50 mM Tris pH 7.5, 500 mM NaCl, and 2% (v/v) glycerol. Protein purification was monitored throughout by SDS-PAGE. AlgJ*_Pp_* and AlgX*_Pp_* proteins were purified as described previously (20, 69).

### NMR structure determination

NMR studies on ^1^H,^15^N,^13^C AlgF ^30-216^ were carried out at a protein concentration of 1 mM in 50 mM phosphate buffer pH 6.8, 10 mM DTT, 2% (v/v) glycerol, 10% D_2_O. To produce uniformly labeled protein, cells were grown in minimal media supplemented with 1 g of ^15^N-NH_4_Cl and 2 g of ^13^C-glucose per litre, and the protein was expressed and purified as described above. The NMR spectra were collected at QANUC on either a Varian INOVA 500 MHz or 800 MHz NMR spectrometer with triple resonance cryoprobes. Backbone resonances were assigned using HNCACB, CBCA(CO)NH, HNCA, and HNCO triple resonance experiments and side-chain resonances were assigned using CCC-TOCSY, HCC-TOCSY and CT-HSQC experiments (31). N- and C-NOESY-HSQC (both aliphatic and aromatic) were used to obtain NOE distance restraints for structural determination purposes. Data were processed with NMRPipe and visualized with NMRDraw (29). Spectral analysis was performed with either NMRView (28, 30) or Analysis by CcpNmr (32, 33). Structures were calculated using CYANA (27) and with the CS-Rosetta server (26), and the models were visualized using PyMOL (The PyMOL Molecular Graphics System, Version 1.2, Schrödinger, LLC). NOE derived models were compared to restraints using the Protein Structure Validation Suite (35) and the Analysis integrated RPF protocol (PyRPF) (32, 33). The CS-Rosetta determined ensembles of the N- and C-terminal domains have been deposited in the PDB with accession codes 6CZT and 6D10 respectively. Relevant NMR data has been deposited in the Biological Magnetic Resonance Bank (BMRB), accession code 30450.

### Structure analysis tools

Inter-residue contacts were determined by CMview (34) and visualized in Excel. Conservation analysis was performed using the ConSurf server with their automatically generated homolog search and multiple sequence alignment (37). Coulombic surface potentials were calculated in ChimeraX (70). Structures were visualized in PyMol and ChimeraX. A tertiary structure comparison search was conducted using the DALI server for the CYANA-determined AlgF N- and C-terminal domains (39).

### Acetylesterase activity assay

Reactions were carried out as previously described (19, 20). Briefly, reactions contained 5 μM of each protein (AlgJ_Pa_ ^79-379^, AlgF_Pa_ ^30-216^, AlgX) in 50 mM sodium HEPES pH 7.6 and 75 mM NaCl at 25 °C and were initiated with the addition of 3-carboxyumbelliferyl acetate (ACC) to 2 mM (dissolved in DMSO at stock concentrations of 20 mM). Deacetylated poly-mannuronate was prepared from *P. aeruginosa* FRD462 as previously described (71) and added at a concentration of 1 mg/mL. The final concentration of DMSO in each reaction was 2% (v/v). Hydrolysis was measured by fluorescence for 20 min with an excitation and emission wavelength of 386 and 446 nm, respectively (72). Reaction rates were calculated using a calibration curve for 7-hydroxycoumarin-3-carboxylic acid, the fluorescent hydrolysis product of ACC. Background hydrolysis rates were measured and subtracted from reaction rates. Assays were carried out in triplicate in 96-well microtitre plates and measured using a SpectraMax M2 microplate reader (Molecular Devices, Sunnyvale, CA). Data analysis was carried out in Prism 7 (Graph Pad) and statistical analyses were performed using an ordinary one-way ANOVA.

### Alginate binding assay

Assays were performed as previously described (20) with a range of synthesized oligomannuronic acid oligomers (73).

### Isothermal titration calorimetry (ITC)

*P. putida* protein constructs were used for ITC analyses. ITC experiments were performed with a MicroCal Auto-ITC200 (Malvern) in a buffer consisting of 50 mM Tris pH 7.8, 150 mM NaCl, 2% (v/v) glycerol, 50 mM arginine, and 50 mM glutamic acid at 20 °C. The Arg/Glu buffer components were used to reach the higher protein concentrations required for ITC experiments (74). ITC mixtures included: 1.86 mM AlgJ (syringe) titrated into 188 μM AlgF (cell) for AlgJ:AlgF; 4.17 mM AlgF (syringe) titrated into 500 μM AlgX (cell) for AlgF:AlgX; 2.97 mM AlgJ (syringe) titrated into 170 μM AlgX (cell) for AlgJ:AlgX. AlgX was placed in the cell due to a lower yield of protein. Runs were performed using 16-20 2.5 μL injections at an interval of 240 s. Data were analyzed using MicroCal Origin ITC Analysis software (Malvern).

### P. aeruginosa strain construction

In-frame, unmarked *algJ* and *algI* gene deletions in *P. aeruginosa* PAO1 Δ*wspF* P_BAD_*alg* (46) were generated using an established allelic replacement protocol (66). Construction of the gene deletion alleles was performed by amplifying flanking regions of the *algJ* or *algI* open reading frames (ORFs) and joining these flanking regions by splicing-by-overlap extension PCR (primers are listed in Table S3). The upstream forward and downstream reverse primers were tailed with EcoRI and HindIII restriction sequences, respectively, and the assembled Δ*algJ* and Δ*algI* alleles were cloned into pEX18Gm. The resulting allelic exchange vectors, pEX18Gm::Δ*algJ* and pEX18Gm::Δ*algI*, were selected on LB agar containing 10 μg/mL gentamicin and verified by Sanger sequencing using the M13F and M13R primers (Table S3).

The deletion alleles encoded by pEX18Gm::Δ*algJ* and pEX18Gm::Δ*algI* were introduced into *P. aeruginosa* PAO1 Δ*wspF* P_BAD_*alg* via biparental mating with the donor strain *E. coli* SM10 (75). Merodiploids were selected on VBMM agar containing 60 μg/mL gentamicin. SacB-mediated counter selection was carried out to select for double cross-over mutations on no-salt lysogeny broth agar containing 15% (w/v) sucrose. Unmarked gene deletions were identified by colony PCR using primers that targeted the outside, flanking regions of *algJ* or *algI* (Table S3). These PCR products were Sanger sequenced using the same primers to confirm the correct deletion.

Construction of strains encoding C-terminally VSV-G-tagged AlgJ, AlgF, and AlgX was performed as above, with the following modifications. The upstream and downstream regions flanking the stop codon of *algX*, *algJ*, and *algF* were amplified using primer pairs whose upstream reverse and downstream forward primers were tailed with complementary sequence encoding the VSV-G peptide sequence (Table S3). The VSV-G sequence was encoded upstream of the stop codon for each gene. The flanking upstream and downstream PCR products were then assembled by splicing-by-overlap extension PCR and cloned into pEX18Gm using SacI and HindIII restriction sites for AlgX and AlgJ, and EcoRI and HindIII restriction sites for AlgF.

### miniTn7 complementation

For gene complementation in *P. aeruginosa*, pUC18T-miniTn7T-Gm, which allows for single-copy chromosomal insertion of genes (76), was modified to allow for arabinose-dependent expression of complemented genes. The *araC*-P_BAD_ promoter from pJJH187 (77) was amplified using the primer pair miniTn7 pBAD F and miniTn7 pBAD R, the latter of which contains flanking sequence encoding SmaI, NotI, PstI, and NcoI sites to generate a multiple cloning site downstream of the *araC*-P_BAD_ promoter (Table 3). The resulting PCR product was cloned into the SacI and HindIII sites of pUC18T-miniTn7T-Gm to generate pUC18T-miniTn7T-Gm-pBAD (Table S2).

The ORF corresponding to *algJ* and *algI* was amplified using the primer pairs AlgJ miniTn7 F + AlgJ miniTn7 R, and AlgI miniTn7 F + AlgI miniTn7 R, respectively, which encode a synthetic ribosome binding site upstream of the start codon (Table S3). The resultant PCR products were cloned into pUC18T-miniTn7T-Gm-pBAD using the NcoI + SacI and NotI + NcoI sites, respectively, selected on LB agar containing 10 μg/mL gentamicin and 100 μg/mL carbenicillin, and confirmed by Sanger sequencing using the miniTn7 SeqF and miniTn7 SeqR primers (Table S3).

Complemented *P. aeruginosa* strains were generated through incorporation of miniTn7 vectors at the neutral *attTn7* site on the *P. aeruginosa* chromosome via electroporation of miniTn7 vectors, along with the helper plasmid pTNS2, as previously described (67). Transposon mutants were selected on LB agar containing 30 μg/ml gentamicin.

### Co-immunoprecipitation (Co-IP)

1 L of LB, containing 0.5% (w/v) L-arabinose and 30 μg/mL gentamicin, was inoculated with a *P. aeruginosa* strain carrying a VSV-G-tagged alginate protein and allowed to grow overnight at 37 °C with shaking. The next day, cells were collected at 5,000 × *g* for 20 min at 4 °C, resuspended in 50 mL of lysis buffer (20 mM Tris pH 8.0, 100 mM NaCl, 1 mM EDTA, 1 mg/mL lysozyme, 100 μg/mL DNase I, 2% (w/v) Triton X-100, 1 SIGMAFAST EDTA-free protease inhibitor cocktail tablet), and rocked for 2 h at 4 °C to allow for cell lysis. The cell lysate was subsequently clarified by centrifugation at 30,000 × *g* for 30 min at 4 °C. A sample of the clarified whole cell lysate was collected before application to the IP resin as a representative example of the input into the experiment. The IP resin (Sigma anti-VSV-G monoclonal antibody-agarose) was prepared by mixing 60 μL of slurry with 10 ml of wash buffer (20 mM Tris pH 8.0, 100 mM NaCl, 2% (w/v) Triton X-100), followed by collection of the IP resin by centrifugation at 100 × *g* for 2 min at 4 °C and removal of the supernatant. The clarified cell lysate was applied to the washed IP resin and incubated at 4 °C for 1 h with agitation. The IP resin was then collected by centrifugation at 100 × *g* for 2 min at 4 °C and the supernatant discarded. The resin was washed four times with 10 ml of wash buffer as above, followed by one wash with 10 mL of detergent-free wash buffer (20 mM Tris pH 8.0, 100 mM NaCl) to remove non-specifically bound protein. Protein was then eluted from the resin by incubation in 110 μL of 0.2 M glycine, pH 2.2, for 15 min at room temperature, followed by collection of the resin by centrifugation at 100 × *g* for 2 min at 4 °C and removal of the supernatant containing eluted protein. The eluate was then neutralized by the addition of 40 μl of 1 M K_2_HPO_4_. Samples of the eluate were analyzed by ESI-MS by the SPARC Biocentre (The Hospital for Sick Children). For Western blot analysis, equal volumes of eluate and 2× Laemmli buffer were mixed, heated at 95 °C for 10 min, and separated by SDS-PAGE followed by Western blot as described below. As a negative control, an IP experiment was also performed using a *P. aeruginosa* strain expressing the corresponding untagged alginate protein.

### Western blot sample preparation and analysis

To analyze protein levels from alginate-overproducing *P. aeruginosa* strains, 5 ml of LB containing 0.5% (w/v) L-arabinose was inoculated with the appropriate strain and allowed to grow overnight at 37 °C with shaking. The next day, culture density was normalized to an OD_600_ = 1 and 1 mL of the resulting culture was centrifuged at 5,000 × *g* for 5 min to pellet cells. The cell pellet was resuspended in 100 μL of 2× Laemmli buffer, heated for 10 min at 95 °C and analyzed by SDS-PAGE followed by Western blot.

For Western blot analysis, a 0.2 μm polyvinylidene difluoride (PVDF) membrane was wetted in methanol and soaked for 5 min in Western transfer buffer (25 mM Tris-HCl, 150 mM glycine, 20% (v/v) methanol) along with the SDS-PAGE gel to be analyzed. Protein was transferred from the SDS-PAGE gel to the PVDF membrane by wet blotting (25 mV, 2 h). The membrane was briefly rinsed in Tris-buffered saline (10 mM Tris-HCl pH 7.5, 150 mM NaCl) containing 0.5% (v/v) Tween-20 (TBS-T) before blocking in 5% (w/v) skim milk powder in TBS-T for 2 h at room temperature with gentle agitation. The membrane was briefly washed again in TBS-T before incubation overnight with primary antibody (1:1,000 α-AlgJ, 1:250 α-AlgF, 1:1,000 α-AlgX, 1:3,000 α-PilP; described below) in TBS-T with 1% (w/v) skim milk powder at 4 °C. The next day, the membrane was washed four times in TBS-T for 5 min each before incubation for 1 h with secondary antibody (1:2,000 dilution of BioRad affinity purified goat α-rabbit IgG conjugated to alkaline phosphatase) in TBS-T with 1% (w/v) skim milk powder. The membrane was then washed three times with TBS-T for 5 min each before development with 5-bromo-4-chloro-3-indolyl phosphate/nitro blue tetrazolium chloride (BioShop ready-to-use BCIP/NBT solution). Developed blots were imaged using a BioRad ChemiDoc imaging system.

### Antibody production and purification

PilP and AlgX antisera were generated and purified as described previously (19, 78). AlgF_Pa_ ^30-216^ and AlgJ ^79-379^ were purified as described above and used to generate antiserum from rabbits via a standard 70 day protocol (Cedarlane Laboratories). The α-AlgF and α-AlgJ antibodies were further purified using a protocol adapted from Salamitou *et al.* (79). Briefly, 200 μg of purified AlgF_Pa_ ^30-216^ or AlgJ_Pa_ ^79-379^ were loaded on a 16% or 12% Tris-HCl polyacrylamide gel, respectively and transferred to a PVDF membrane. The membrane was stained with Ponceau S and the band corresponding to AlgF_Pa_ ^30-216^ or AlgJ_Pa_ ^79-379^ was cut out and blocked using PBS pH 7.0 with 0.1% (w/v) Tween 20 and 5% (w/v) skim milk powder for 1 h. The membrane was then incubated with α-AlgF or α-AlgJ antisera overnight at 4 °C followed by incubation at room temperature for 2 h. After washing in PBS, α-AlgF or α-AlgJ antibodies were eluted from the membrane by incubation in 700 μL of 0.2 M glycine pH 2.2 for 15 min, followed by neutralization with 300 μL of 1 M K_2_HPO_4_. The purified antibodies were dialyzed overnight against PBS, mixed 1:1 with glycerol, and stored at −20 °C. α-AlgF antibodies were used at a dilution of 1:250, α-AlgJ antibodies were used at a dilution of 1:1,000, α-AlgX antibodies were used at a dilution of 1:1,000, and anti-PilP antibodies were used at a dilution of 1:3,000.

### AlphaFold modelling

AlphaFold predictions were run using ColabFold v 1.5.2 with default parameters (i.e. alphafold2_multimer_v3). No template information was used. and the multiple sequence alignment options chosen were mmseq2_uniref_env and unpaired_paired. Five structures for each complex were predicted and used without relaxation using Amber. The sequences for *P. putida* were retrieve from UniProt accession numbers (AlgI: Q88ND2 AlgJ: Q88ND3; AlgF: Q88ND4; AlgX: Q88ND0; AlgK: Q88NC7) were used either as is (AlgI, AlgJ) or with their signal sequences (as determined by SignalP (68)) removed (AlgF, AlgX, and AlgK).

### Software support

The majority of software programs used in this report were configured and supported by the SBGrid consortium (80).

## Acknowledgements

Thanks to Patrick Yip, Gaelen Moore, Jeffrey Lynham, Tyler Ricer, and Robert Vernon for technical support and to Perrin Baker for helpful discussion. Thanks to Brookhaven National Laboratory NSLS-II 16-ID beamline staff for technical support and helpful discussion. The acetylesterase pseudosubstrate 3-carboxyumbelliferyl acetate (ACC) was generously donated by Mark Nitz. This research was supported by the Canadian Institutes of Health Research (#MOP-13337, PJT-153322, and FDN-154327). GBW was supported by graduate scholarships from NSERC Canada Graduate Scholarships and Cystic Fibrosis Canada. AAG was supported by graduate scholarship from The SickKids Foundation (RESTRACOMP) and Cystic Fibrosis Canada. YEL was supported by the GlycoNet Summer Awards Program for Undergraduates. JTW was supported by a Postdoctoral Fellowship from NSERC. PLH is the recipient of a Tier I Canada Research Chair. The studies described herein used equipment in the Structural and Biophysical Core Facility at The Hospital for Sick Children, which is funded in part by the Canadian Foundation for Innovation.

## Author contributions

A.A.G., K.E.L., S.D.T., G.B.W., L.M.R., J.T.W., and P.L.H designed the research; K.E.L., S.D.T., G.B.W., A.A.G., Y.E.L., L.M.R., J.T.W., S.J.C., P.A.C., M.T.C.W., and E.N.K. performed the research; K.E.L., S.D.T., G.B.W., A.A.G., Y.E.L., L.M.R., J.T.W., S.J.C., M.T.C.W., E.N.K., J.S.K., and J.D.C.C. analyzed the data; K.E.L., S.D.T., G.B.W., L.M.R., A.A.G., and P.L.H. wrote the paper. All authors provided feedback and approved the final manuscript.

## Additional information

### Data Deposition

The CS-Rosetta determined ensembles of the N- and C-terminal domains have been deposited in the PDB with accession codes 6CZT and 6D10, respectively. Relevant NMR data has been deposited to the BMRB under accession code 30450.

