## Supplementary material for "*Pseudomonas aeruginosa* AlgF is a protein-protein interaction mediator required for acetylation of the alginate exopolysaccharide": SI figures

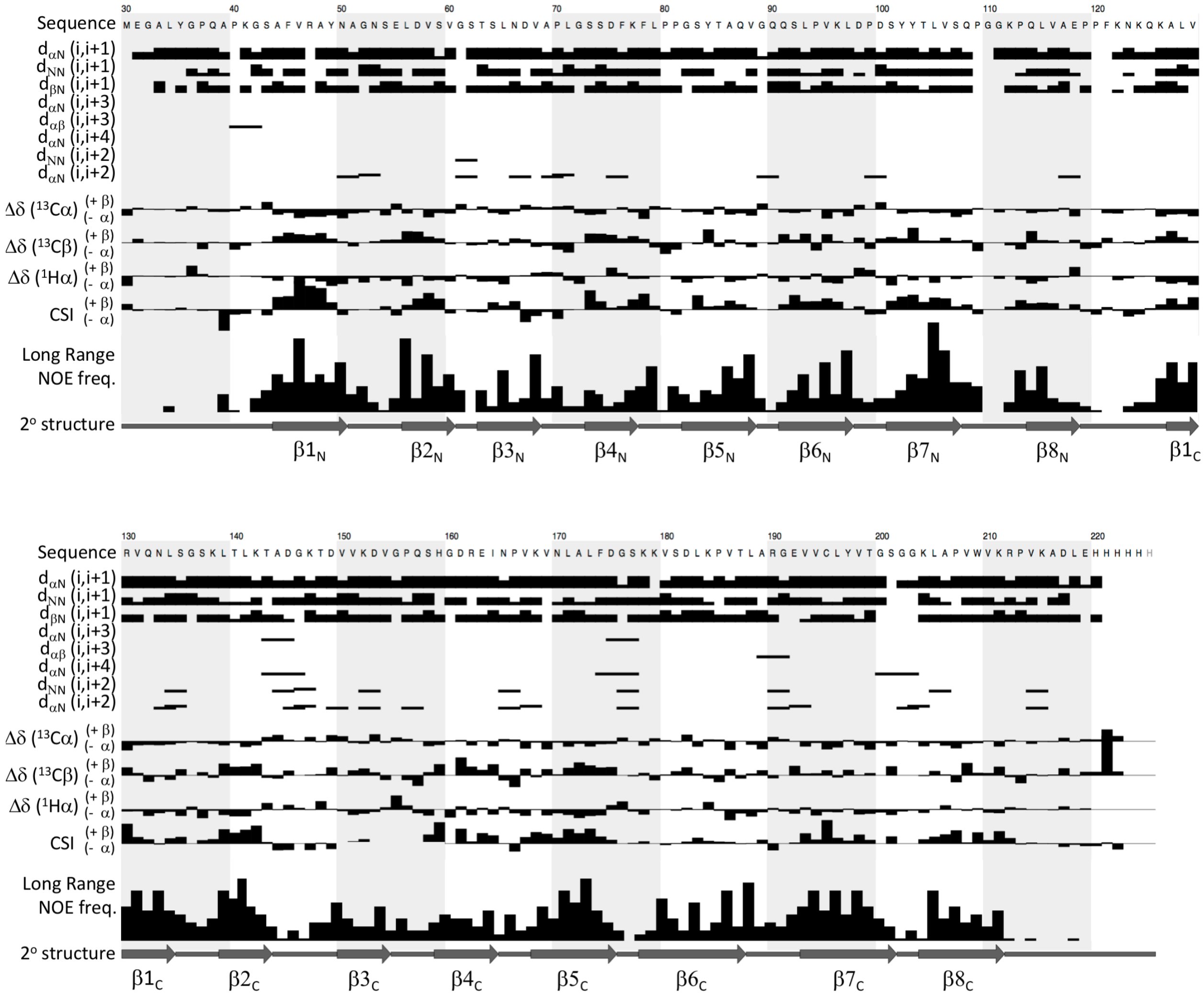


**Figure S1. Chemical Shift and NOE restraint analysis.** The sequence and amino acid numbering of the $AlgF_{Pa}^{30\text{-}216}$ construct used in the NMR experiments is shown in the top row. The next 8 rows are NOE assignments (short range NOEs between adjacent a and amide protons (labeled as i-i+1) and medium range NOEs between α, β, and amide protons (labeled as i-i+2, i-i+3, i-i+4)). The short-range NOEs show the completeness of the backbone assignments, while the medium range NOE pattern indicates α-helical or β-turn structures. The next 4 rows are the chemical shift index (CSI) analysis. The deviance in chemical shift from random coil values (Δδ) for $C^{\alpha}$, $C^{\beta}$, and $H^{\alpha}$ resonances are shown as a bar graph with the overall average CSI in the final row. Positive Δδ values are found in β-strand structures while negative Δδ values are found in α-helical structures. Next is the frequency of long-range NOE assignment (between residues more than 4 residues away in 1° sequence). The frequency of long-range assignments is lowest at the termini and loops/interdomain region. The final row is the 2° structure as observed in the NMR chemical shift data and calculated models. The secondary structural elements are labeled with a subscript N and C for the N- and C-terminal domains, respectively.

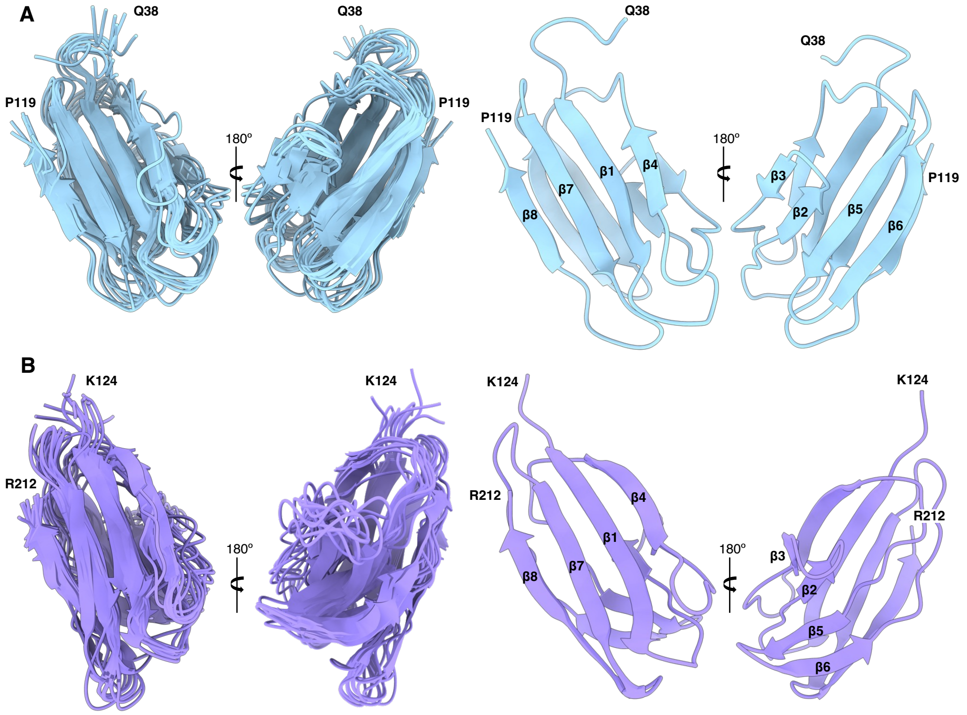


**Figure S2: CS-Rosetta ensembles for the AlgF$_{Pa}$ N- and C-terminal domains.** (A) Left panel shows the superposition of the 10 models in the N-terminal domain ensemble (RMSD 1.38 Å over 304 backbone atoms). Right panel is the lowest energy model in the same orientation as the ensemble. Secondary structure elements are labelled for each face. In all cases the terminal residues are labeled. (B) As in (A) but for the C-terminal domain. The ensemble of 10 models has an RMSD of 1.59 Å over 340 backbone atoms.

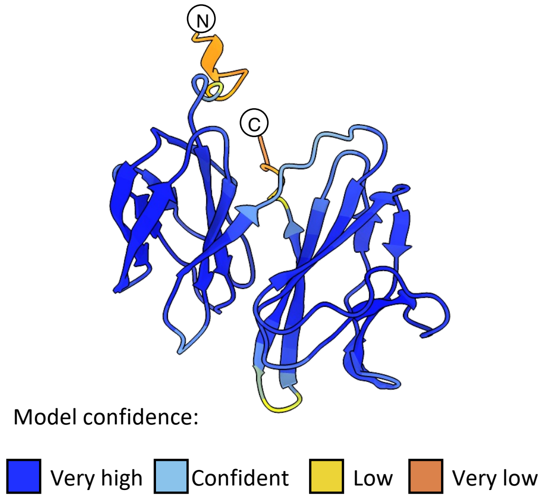


**Figure S3: Confidence score of the AlphaFold2 predicted AlgF$_{Pa}$ model.** Regions are coloured by confidence level.

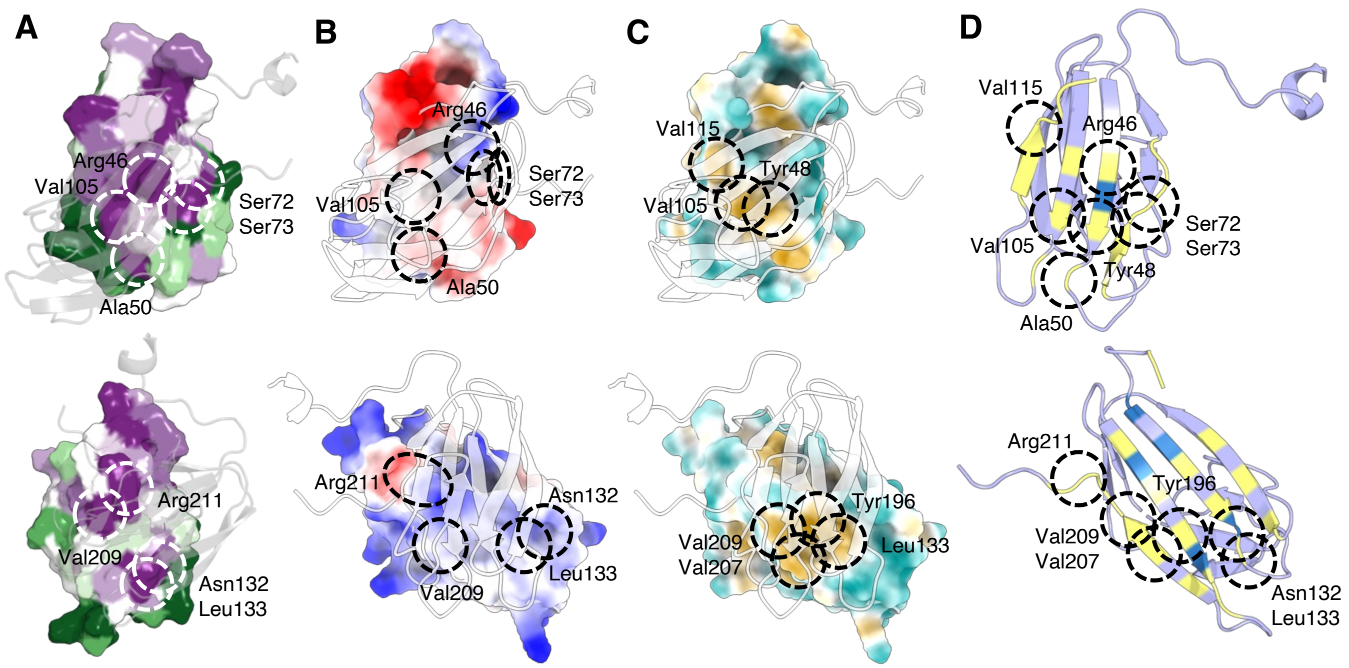


**Figure S4: Conservation, surface electrostatics, hydrophobicity and buried surface analyses support the compact structure predicted by AlphaFold for AlgF$_{Pa}$.** (A) AlgF$_{Pa}$ N-terminal domain surface representation demonstrating conservation superimposed with the AlgF$_{Pa}$ AlphaFold model in grey (top). AlgF$_{Pa}$ C-terminal domain surface representation demonstrating conservation superimposed with the AlgF$_{Pa}$ AlphaFold model in grey (bottom). Surface representation colored by level of conservation (green variable to purple highly conserved). White dashed circles indicate amino acid residues of interest. (B) AlgF$_{Pa}$ N-terminal domain surface representation demonstrating surface electrostatics superimposed with the AlgF$_{Pa}$ AlphaFold model in grey (top). AlgF$_{Pa}$ C-terminal domain surface representation demonstrating surface electrostatics superimposed with the AlgF$_{Pa}$ AlphaFold model in grey (bottom). The coulombic potential range from -10 (red) to 10 (blue) kcal/(mol · $e$). Black dashed circles indicate amino acid residues of interest. (C) AlgF$_{Pa}$ N-terminal domain surface representation demonstrating surface hydrophobicity superimposed with the AlgF$_{Pa}$ AlphaFold model in grey (top). AlgF$_{Pa}$ C-terminal domain surface representation demonstrating surface electrostatics superimposed with the AlgF$_{Pa}$ AlphaFold model in grey (bottom). Hydrophobicity ranging from hydrophobic (yellow) to hydrophilic (teal). Black dashed circles indicate amino acid residues of interest. (D) AlgF$_{Pa}$ N-terminal domain (top) and AlgF$_{Pa}$ C-terminal domain (bottom) depicting the exposed and buried residues as calculated by PISA. Solvent exposed residues (light blue), buried residues (yellow), and solvent inaccessible residues (dark blue). Black dashed circles indicate amino acid residues of interest.

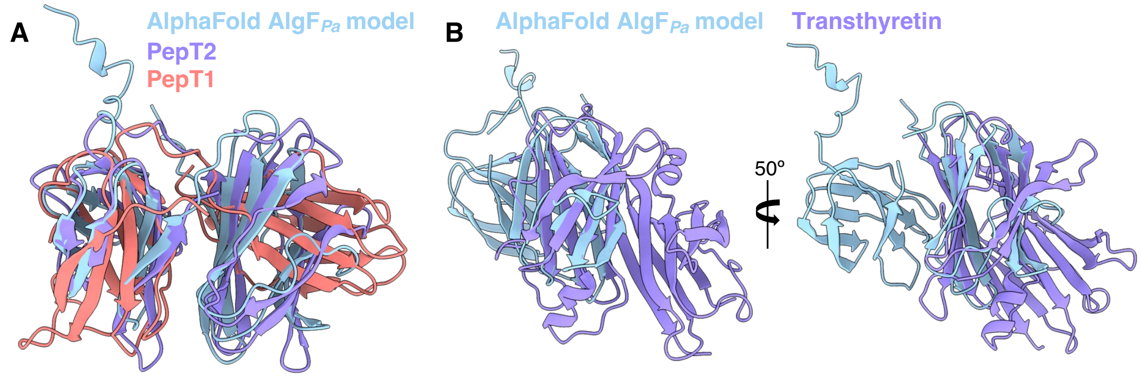


**Figure S5: Structural superimposition of the AlgF$_{Pa}$ AlphaFold model with select DALI search results.** (A) Structural superimposition of the AlgF$_{Pa}$ AlphaFold model (light blue) with PepT2 (PDB: 5A9H) (purple) and PepT1 (PDB: 5A9I) (red). (B) Structural superimposition of the AlgF$_{Pa}$ AlphaFold model (light blue) with transthyretin (PDB: 6R66) (purple).

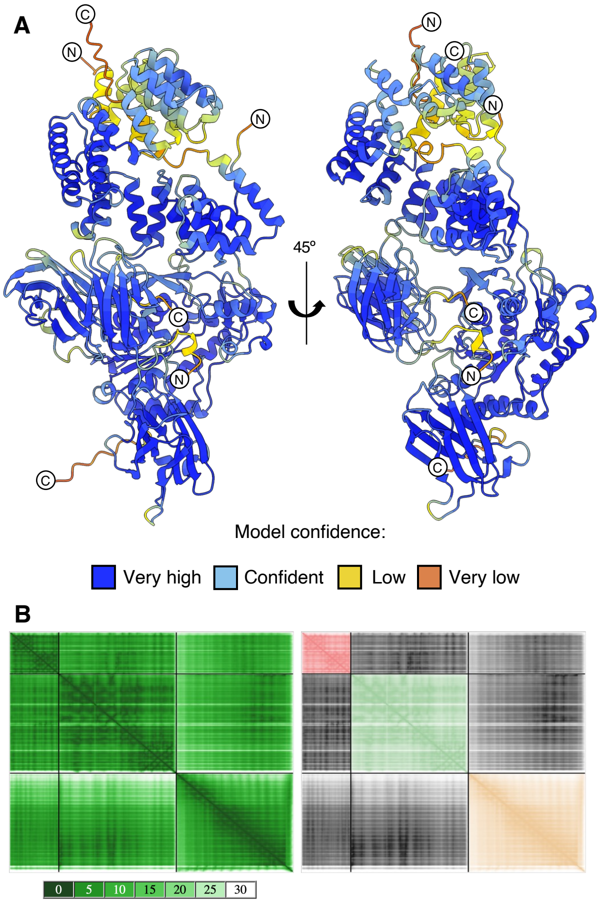


**Figure S6: Confidence score and PAE plot of the AlphaFold2 predicted AlgFXK$_{Pp}$ model.** (A) The structures are coloured by confidence level. (B) Residue-residue predicted aligned error (PAE) values for the top ranked AlphaFold2 model of the AlgKXF$_{Pa}$ complex. Left: The PAE values give a distance error for every pair of residues and is an estimate of the positional error at residue x when the predicted and true structures are aligned. The values range from 0-30 Å, (dark green to white). If the relative position of two domains is confidently predicted the PAE value between two residues will be less than 5Å. Right: The PAE plot is coloured by protein as depicted in Figure 7: AlgF (pink), AlgX (green) and AlgF (gold).

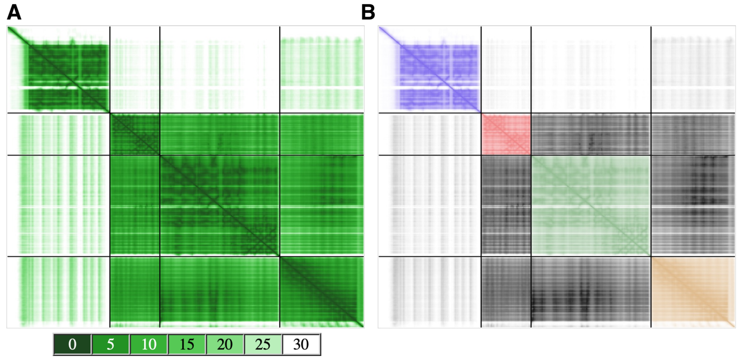


**Figure S7: PAE plot of the AlphaFold2 predicted AlgJFXK*Pp* model.** (A) Residue-residue predicted aligned error (PAE) values for the predicted by AlphaFold2 for the AlgJFX*Pp* and C-terminal region of AlgK*Pa*. Plot is coloured by error value in Å from 0-30Å (dark green to white. (B) The PAE plot coloured by protein as depicted in Figure 7: AlgF (pink), AlgX (green) and AlgK (gold) with AlgJ in purple. The PAE values between AlgJ and the rest of the protein complex are > 25Å.

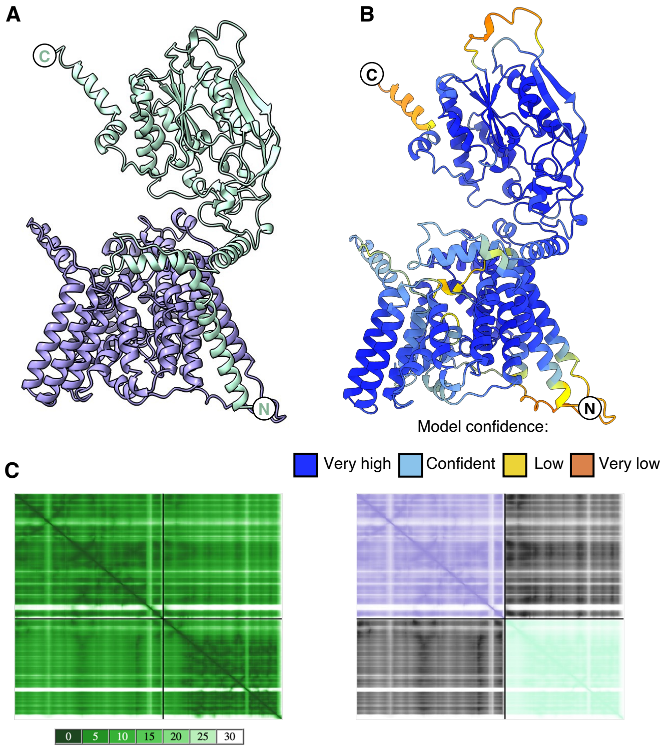


**Figure S8: Confidence score and PAE plot of the AlphaFold2 predicted AlgIJ$_{Pp}$ model.** (A) Predicted AlphaFold2 model of the AlgIJ$_{Pp}$ complex. AlgI$_{Pp}$ (purple), AlgJ$_{Pp}$ (cyan). (B) AlgIJ$_{Pp}$ structures are coloured by confidence level. (C) PEA plot coloured by error value in Å from 0-30Å (dark green to white) (left) and by protein AlgI (purple) and AlgJ (cyan). N- and C-termini of AlgJ$_{Pp}$ are indicated. N- and C-termini of AlgI$_{Pp}$ are not visible in the presented orientation.

**Table S1: Apparent association constants ($K_a$) for AlgF*Pp* binding to poly-mannuronate polymers at 25ºC and pH 7.0 determined by direct ESI-MS.**

| Ligand | Apparent $K_a$ ($M^{-1}$) |
|---|---|
| $ManA_4$ | <500 |
| $ManA_5$ | <500 |
| $ManA_6$ | <500 |
| $ManA_7$ | <500 |
| $ManA_8$ | $<10^3$ |
| $ManA_9$ | $<10^3$ |
| $ManA_{10}$ | $<10^3$ |
| $ManA_{11}$ | $<10^3$ |
| $ManA_{12}$ | $<10^3$ |

**Table S2: Bacterial strains and plasmids used in this study**

| Strain | Description | Reference |
|---|---|---|
| ***E. coli*** | | |
| TOP10 | Cloning strain; F⁻ *mcrA*Δ(*mrr-hsdRMS-mcrBC*) Φ80*lacZ* ΔM15 Δ*lacX*74 *recA*1 *araD*139Δ(*ara leu*) 7697 *galU galK rpsL endA*1 *nupG*, $Str^R$ | Invitrogen |
| DH5α | Cloning strain; F⁻ Φ80*lacZ*ΔM15 Δ(*lacZYA-argF*) U169 *recA1 endA1 hsdR17* ($r_K^-$, $m_K^+$) *phoA supE44* λ⁻ *thi*-1 *gyrA96 relA1* | Invitrogen |
| SM10 | Bi-parental mating strain; *thi thr leu tonA lacY supE recA*::RP4-2-Tc::Mu $K_m$ λ*pir*, $Kan^R$, $Tet^R$ | (1) |
| BL21CodonPlus™ (DE3)-RP | Protein expression strain; F⁻ *ompT hsdS*($r_B^-$ $m_B^-$)*dcm*⁺ $Tet^R$ *gal*λ (DE3) *endA*Hte *metA*::Tn5($Kan^R$) [*argU ileY leuW* $Cam^R$] | Stratagene |
| ***P. aeruginosa*** | | |
| PAO1 | Wild-type strain | M. R. Parsek |
| PAO1 Δ*wspF* $P_{BAD}$*alg* | PAO1 Δ*wspF* (in-frame); *araC*-$P_{BAD}$ inserted upstream of *algD* operon | (2) |
| GBW7 | PAO1 Δ*wspF* $P_{BAD}$*alg* Δ*algJ* | This study |
| GBW8 | PAO1 Δ*wspF* $P_{BAD}$*alg* Δ*algI* | This study |
| GBW9 | GBW7 *attTn7*::miniTn7T-Gm::*araC*-$P_{BAD}$::*algJ*, $Gen^R$ | This study |
| GBW10 | GBW8 *attTn7*::miniTn7T-Gm::*araC*-$P_{BAD}$::*algI*, $Gen^R$ | This study |
| GBW11 | PAO1 Δ*wspF* $P_{BAD}$*alg* *algX*$^{C\text{-}VSV\text{-}G}$ | This study |
| GBW12 | PAO1 Δ*wspF* $P_{BAD}$*alg* *algJ*$^{C\text{-}VSV\text{-}G}$ | This study |
| GBW13 | PAO1 Δ*wspF* $P_{BAD}$*alg* *algF*$^{C\text{-}VSV\text{-}G}$ | This study |

| Plasmid | Description | Reference |
|---|---|---|
| **Recombinant protein expression** | | |
| pET24b | IPTG-inducible expression vector encoding C-terminal hexahistidine tag, $Kan^R$ | Novagen |
| pET28a | IPTG-inducible expression vector encoding N-terminal hexahistidine tag and thrombin cleavage site, $Kan^R$ | Novagen |

| | | |
|---|---|---|
| pET24b::AlgF$_{Pa}^{30-216}$ | pET24b with *P. aeruginosa* PAO1 *algF* corresponding to residues 30-216 cloned between the NdeI and XhoI sites; Kan$^R$ | This study |
| pET24b::AlgX$_{Pa}$ | pET24b with *P. aeruginosa* PAO1 *algX* cloned between the NdeI and XhoI sites,codon-optimized for expression in *E. coli*; Kan$^R$ | This study |
| pET28a::AlgF$_{Pp}^{29-215}$ | pET28a with *P. putida* KT2440 *algF* corresponding to residues 29-215 cloned between the NdeI and XhoI sites; Kan$^R$ | This study |
| pET28a::AlgF$_{Pp}^{29-120}$ | pET28a with *P. putida* KT2440 *algF* corresponding to residues 29-120 cloned between the NdeI and XhoI sites; Kan$^R$ | This study |
| pET28a::AlgF$_{Pp}^{122-215}$ | pET28a with *P. putida* KT2440 *algF* corresponding to residues 122-215 cloned between the NdeI and XhoI sites; Kan$^R$ | This study |
| pET28a::AlgJ$_{Pa}^{79-379}$ | pET28a with *P. aeruginosa* PAO1 *algJ* corresponding to residues 79-379 cloned between the NdeI and XhoI sites; Kan$^R$ | (3) |
| pET28a::AlgJ$_{Pp}^{75-370}$ | pET28a with *P. putida* KT2440 *algJ* corresponding to residues 75-370 cloned between the NdeI and XhoI sites; Kan$^R$ | (3) |
| pET24b::AlgX$_{Pp}$ | pET24b with *P. putida* KT2440 *algX* cloned between the NdeI and XhoI sites, codon-optimized for expression in *E. coli*; Kan$^R$ | This study |
| **Allelic exchange** | | |
| pEX18Gm | Suicide vector for allelic exchange in *P. aeruginosa*, encodes SacB, Gen$^R$ | (1) |
| pEX18Gm::Δ*algJ* | pEX18Gm with *P. aeruginosa* PAO1 Δ*algJ* cloned between the EcoRI and HindIII sites, Gen$^R$ | This study |
| pEX18Gm::Δ*algI* | pEX18Gm with *P. aeruginosa* PAO1 Δ*algI* cloned between the EcoRI and HindIII sites, Gen$^R$ | This study |
| pEX18Gm::*algX*$^{C\text{-}VSV\text{-}G}$ | pEX18Gm containing *P. aeruginosa* PAO1 *algX*$^{C\text{-}VSV\text{-}G}$ cloned between the SacI and HindII sites, Gen$^R$ | This study |
| pEX18Gm::*algJ*$^{C\text{-}VSV\text{-}G}$ | pEX18Gm containing *P. aeruginosa* PAO1 *algJ*$^{C\text{-}VSV\text{-}G}$ cloned between the SacI and HindII sites, Gen$^R$ | This study |
| pEX18Gm::*algF*$^{C\text{-}VSV\text{-}G}$ | pEX18Gm containing *P. aeruginosa* PAO1 *algF*$^{C\text{-}VSV\text{-}G}$ cloned between the EcoRI and HindII sites, Gen$^R$ | This study |
| **Complementation analysis** | | |

| | | |
|---|---|---|
| pUC18T-miniTn7T-Gm | *aacC1* on miniTn7-based vector with transcriptional terminators at the right end of the Tn7 transposon; $Amp^R$, $Gen^R$ | (4) |
| pUC18T-miniTn7T-Gm-pBAD | pUC18T-miniTn7T-Gm containing *araC*-$P_{BAD}$ and a downstream MCS (SmaI-NotI-PstI-NcoI) cloned between the HindIII and SacI sites; $Amp^R$, $Gen^R$ | This study |
| pTNS2 | Helper plasmid encoding *tnsABCD*, $Amp^R$ | (5) |
| p-miniTn7-AlgJ | pUC18T-miniTn7T-Gm-pBAD with *P. aeruginosa* PAO1 *algJ* fused to an upstream synthetic ribosome binding site, cloned between the NcoI and SacI sites; $Amp^R$, $Gen^R$ | This study |
| p-miniTn7-AlgI | pUC18T-miniTn7T-Gm-pBAD with *P. aeruginosa* PAO1 *algI* fused to an upstream synthetic ribosome binding site, cloned between the NotI and NcoI sites; $Amp^R$, $Gen^R$ | This study |

**Table S3: Primers used in this study**

| Name | Sequence |
|---|---|
| **Recombinant protein production** | |
| AlgF$_{Pa}$ 5’ | GCG **CCA TGG** AGG GCG CCC TG |
| AlgF$_{Pa}$ 3’ | GCG **CTC GAG** ATC CGC CTT CAC CGG |
| AlgF$_{Pp}$ 5’ | GTT **CAT ATG** GAC GCC GCA CTG TAT GGC CC |
| AlgF$_{Pp}$ 3’ | GTT **CTC GAG** TCA GTT GCT AGC CAC GGG GC |
| **Allelic exchange** | |
| ΔalgJ upF | GGG **GAA TTC** TGG CGC ATG TAC GAA GCC AT |
| ΔalgJ upR | *TCA ATC GTT GCG GCT GGC GGC* GCG GGA AAT GCT CTG TGT CAT |
| ΔalgJ downF | GCC GCC AGC CGC AAC GAT TGA |
| ΔalgJ downR | GAG **AAG CTT** GCT CGG CCA CCA GCT GCG GC |
| ΔalgI upF | GGG **GAA TTC** AGA AGA CCG GCG AGG ACC AGG AC |
| ΔalgI upR | *TCA GAA CTG GAA GTA GAG GAA TGG CGA* CAG GAA CAG GAA CAG GAA CAC GTT TGA |
| ΔalgI downF | TCG CCA TTC CTC TAC TTC CAG TTC TGA |
| ΔalgI downR | CAT **AAG CTT** TCA CCG CGA GCA CCA GCT TGA C |
| AlgX C-VSV-G upF | GGT **GAG CTC** CAT CTG GGA ATT CGC CAC CC |
| AlgX C-VSV-G upR | *TTT TCC TAA TCT ATT CAT TTC AAT ATC TGT ATA* CCT CCC GGC CAC CGA CTG |
| AlgX C-VSV-G downF | *TAT ACA GAT ATT GAA ATG AAT AGA TTA GGA AAA* TAA ACG ATG AAA ACG TCC CAC C |
| AlgX C-VSV-G downR | GGG **AAG CTT** GCT GCC CAG CGC CCA TTT G |
| AlgJ C-VSV-G upF | GGG **GAG CTC** AAG GGC GAC CTG CTCAGC TT |
| AlgJ C-VSV-G upR | *TTT TCC TAA TCT ATT CAT TTC AAT ATC TGT ATA* ATC GT TGC GGC TGG CGG C |
| AlgJ C-VSV-G downF | *TAT ACA GAT ATT GAA ATG AAT AGA TTA GGA AAA* TGA ACG AAC ACA GCC AAG GCT |
| AlgJ C-VSV-G downR | GGG **AAG CTT** GAG GTT CTG CAC CCG TAC CA |

| | |
|---|---|
| AlgF C-VSV-G upF | GGG **GAA TTC** AGC ACC AGC CTG AAC GAC GT |
| AlgF C-VSV-G upR | *TTT TCC TAA TCT ATT CAT TTC AAT ATC TGT ATA* GTC CGC CTT CAC CGG GCG |
| AlgF C-VSV-G downF | *TAT ACA GAT ATT GAA ATG AAT AGA TTA GGA AAA* TGA GAC CGA CCG GGA GAG GC |
| AlgF C-VSV-G downR | GGG **AAG CTT** GAA GGG TTC GAG GAG GAT CG |
| **miniTn7 complementation** | |
| miniTn7 pBAD F | GGG **AAG CTT** TTA TGA CAA CTT GAC GGC TA |
| miniTn7 pBAD R | GGG **GAG CTC** CCA TGG CTG CAG GCG GCC GCC CCG GGC AAA AAA ACG GGT ATG GAG AAA CAG TA |
| AlgJ miniTn7 F | A GT**C CAT GG**G AGG AGG ATA TTC ATG ACA CAG AGC ATT TCC CGC CCC |
| AlgJ miniTn7 R | ACG **GAG CTC** TCA ATC GTT GCG GCT GGC GGC C |
| AlgI miniTn7 F | TAT **GCG GCC GC**G AGG AGG ATA TTC ATG GTC TTT TCT TCA AAC GTG TTC CTG |
| AlgI miniTn7 R | ACG **CCA TGG** TCA GAA CTG GAA GTA GAG GAA TGG CG |
| **Sequencing** | |
| M13F | GTA AAA CGA CGG CCA GT |
| M13R | AAC AGC TAT GAC CAT G |
| AlgJ SeqF | TTC CTC CTG GTG GTG ATC GG |
| AlgJ SeqR | CTT CAG GGT CAG CTT CGA GC |
| AlgI SeqF | CTG GCC GAG CGG GTG ATG AA |
| AlgI SeqR | GTA GAG GCT GTC GTG CAG CG |
| miniTn7 SeqF | GCG GAT CCT ACC TGA CGC TT |
| miniTn7 SeqR | AAC TTC AGA GCG CTT TTG AA |
